# Effects of cortical thickness, volume, and memory performance on age differences in neural reinstatement of scene information

**DOI:** 10.1101/2025.02.06.636759

**Authors:** Joshua M. Olivier, Sabina Srokova, Paul F. Hill, Michael D. Rugg

**Affiliations:** Center for Vital Longevity and School of Behavioral and Brain Sciences, University of Texas at Dallas 1600 Viceroy Drive, Ste. 800 Dallas, TX 75235; Department of Psychology and Evelyn F. McKnight Brain Institute, University of Arizona Tucson, AZ 85721

**Keywords:** episodic memory, fMRI, neural selectivity, neural broadening, structural MRI

## Abstract

The strength of neural reinstatement, a correlate of episodic memory retrieval, reportedly reflects the amount and fidelity of mnemonic content and is weaker in older than younger adults, especially for scene memoranda. Based on evidence that age-related declines in cortical thickness and volume contribute to age-related cognitive decline, we analyzed fMRI data acquired from healthy young and older adults to examine relationships between cortical thickness, cortical volume, age, and scene-related reinstatement in three cortical regions implicated in scene processing: the parahippocampal place area (PPA), medial place area (MPA), and occipital place area (OPA). A ‘reinstatement index’ was estimated from fMRI data collected during tests of source memory for scene images, and multiple regression analyses were employed to examine the effects of the variables of interest on scene reinstatement. There were robust age differences in reinstatement, cortical thickness, and cortical volume. In the PPA and MPA, cortical volume fully mediated the effects of age on reinstatement. Additionally, PPA reinstatement strength predicted source memory performance independently of cortical volume or age. These findings suggest that age differences in scene reinstatement are mediated by cortical volume and that memory performance and cortical volume are associated with unique components of variance in reinstatement strength.

---

Episodic memory, the ability to recollect information about previously experienced events, declines with increasing age (Nilsson 2003; Nyberg et al. 2012). Neuroimaging research has revealed robust evidence for age differences in a variety of functional correlates of episodic memory encoding and retrieval (e.g., Cabeza et al. 2002; Davis et al. 2008; Park et al. 2013; Srokova et al. 2022; Schott et al. 2023). One such correlate is neural reinstatement—retrieval-related neural activity that partially overlaps with the activity elicited during encoding (for reviews, see Danker and Anderson 2010; Rissman and Wagner 2012; Rugg et al. 2015). It has been reported that stronger reinstatement positively covaries with memory performance (Johnson et al. 2009; Gordon et al. 2014; Thakral et al. 2015; Trelle et al. 2020; Srokova et al. 2021), suggesting that the strength of reinstatement reflects the amount and quality of the retrieved mnemonic content. Additionally, it has sometimes been reported that the strength of neural reinstatement varies with age, with older adults showing weaker effects than young adults (e.g., McDonough et al. 2014; St-Laurent et al. 2014; Johnson et al. 2015; Abdulrahman et al. 2017; Folville et al. 2020; Trelle et al. 2020; for opposing results, see Dulas and Duarte 2012; Wang et al. 2016; Thakral et al. 2017; St-Laurent and Buchsbaum 2019; Thakral et al. 2019). In Hill et al. (2021), a study that examined neural reinstatement of face and scene images, age differences were only evident for scene reinstatement (see also Trelle et al. 2020). In keeping with this finding, based on a critical review of the literature, Rugg and Srokova (2024) concluded that the most secure evidence for age-related attenuation of reinstatement effects comes from studies that employed scene images as the critical memoranda. Here, we address the question of whether these age differences are associated with corresponding differences in brain structural metrics. We also examine whether reinstatement is associated across participants with memory performance and whether any such relationship is moderated by age.

Age differences in structural and functional brain metrics have primarily been studied separately, despite the extensive cross-sectional and longitudinal evidence indicating that structural metrics such as cortical volume and thickness positively covary with cognitive abilities in older samples (Karama et al. 2014; Salthouse et al. 2015; Sun et al. 2016; Kranz et al. 2018; de Chastelaine et al. 2019; de Chastelaine et al. 2023). In a recent study, it was reported that age differences in gray matter volume mediated corresponding differences in fMRI ‘subsequent memory effects,’ a functional correlate of episodic memory encoding (Kizilirmak et al. 2024). By contrast, other studies examining relationships between structural and functional measures have reported little evidence for a mediating effect of structure (e.g., Boller et al. 2017; Hou et al. 2020; Hou et al. 2021; Kidwai et al. 2025). Overall, however, the extant literature is limited, and the circumstances under which correlations between brain function and structure can be identified remain unclear.

Motivated by this gap in knowledge, we examined the relationship between neural reinstatement of scene information, cortical thickness, and cortical volume in samples of young and older adults. We combined two independent datasets that comprised a total of 42 young and 40 older adults as they retrieved word-image pairs while undergoing fMRI. The combined dataset provided a substantial boost in statistical power relative to that associated with separate analyses of the two datasets. Our analyses focused on scene reinstatement effects in three regions that have been heavily implicated in visual scene processing: the parahippocampal place area (PPA), medial place area (MPA), and occipital place area (OPA) (Epstein and Baker 2019)—with the aim of examining the effects of age on scene reinstatement and the mediating effect, if any, of the structural metrics of cortical thickness and volume.

## Materials and Methods

The present report examines the relationship between cortical thickness and volume and scene reinstatement by combining data from two independent, previously reported studies (Hill et al. 2021; Srokova et al. 2021). The experimental procedures and behavioral analyses for each study were detailed in those prior reports and are briefly summarized below for the convenience of the reader. Note that in Hill et al. (2021), neural reinstatement was indexed by multi-voxel pattern analysis rather than the univariate metric employed in the present study, and the researchers did not differentiate between the PPA and MPA. In Srokova et al. (2021), unlike here, reinstatement was estimated for all correctly recognized test items regardless of the accuracy of the subsequent source memory judgment. Moreover, neither study reported brain structural metrics. Thus, none of the analyses of the functional and structural data reported below have been described previously.

### Participants and Neuropsychological Testing

The present report describes analyses performed on 42 younger and 40 older adults across two independent studies. All participants were recruited from the Dallas metropolitan area. All were right-handed, had normal or corrected-to-normal vision, and were fluent English speakers before age 5. Potential participants were excluded if they reported a history of neurological or cardiovascular disease, diabetes, substance abuse, or usage of medication affecting the central nervous system. All participants gave informed consent in accordance with the University of Texas at Dallas and University of Texas Southwestern Medical Center Institutional Review Boards, which approved the studies (approval numbers IRB-24-277 and STU 092010-226, respectively) and were compensated at a rate of $30 per hour. Demographic data for the combined participant sample can be found in Table 1.

**Table 1:** Demographic data for participant sample by study (M ± SD)

|  |  | Young adults | Older adults | Group difference (p) |
| --- | --- | --- | --- | --- |
| **Study 1** | *N* | 24 | 22 |  |
|  | Sex (F/M) | 15/9 | 13/9 | 1.00<sup>†</sup> |
| | Age (years)<sup>‡</sup> | 22.42 ± 3.24 | 69.95 ± 3.36 | |
| | Years of Education<sup>‡</sup> | 15.46 ± 2.65 | 16.77 ± 2.47 | .089 |
| **Study 2** | *N* | 18 | 18 |  |
|  | Sex (F/M) | 10/8 | 7/11 | .504<sup>†</sup> |
| | Age (years)<sup>‡</sup> | 24.39 ± 3.55 | 68.17 ± 2.60 | |
| | Years of Education<sup>‡</sup> | 16.00 ± 1.71 | 17.61 ± 2.70 | **.041\*** |

<sup>†</sup>Result from a $\chi^2$ test with Yates' continuity correction.
<sup>‡</sup>Result from one-way ANOVA revealed no significant differences between studies (Age: $p = .830$ ; Years of Education: $p = .203$ ).

Participants in both studies undertook an extensive neuropsychological test battery, including the Mini-Mental State Examination (MMSE), the California Verbal Learning Test-II (CVLT; Delis et al. 2000), Wechsler Logical Memory (Tests 1 and 2; Wechsler 2009a), Trail Making (Tests A and B; Reitan and Wolfson 1985), the Symbol Digit Modalities test (SDMT; Smith 1982), the F-A-S subtest of the Neurosensory Center Comprehensive Evaluation for Aphasia (Spreen and Benton 1977), the Wechsler Adult Intelligence Scale-Revised (forward and backward digit span subtests; Wechsler 1981), Category Fluency test (Benton 1968), Raven’s Progressive Matrices (List 1; Raven et al. 2000), and either the Wechsler Test of Adult Reading (WTAR; Wechsler 2001) or the Wechsler Test of Premorbid Functioning (Wechsler 2009b). Participants were excluded if their performance was greater than 1.5 standard deviations below age norms for one or more memory-based tests or two or more non-memory tests. Participants with scores of less than 27 on the MMSE were also excluded. For a description of the neuropsychological test battery scores in each study, see Hill et al. (2021) and Srokova et al. (2021).

**Study 1:** 27 young and 33 older adults were recruited to participate in Study 1. Three young and nine older participants were excluded from the initial sample due to inadequate memory performance, incidental MRI findings, technical difficulties, or voluntary withdrawal from the study. Two additional older participants were excluded from the analyses reported here due to the poor quality of their T1-weighted structural images caused by excessive movement. Thus, the final sample for Study 1 included 24 young (age range: 18–28 years) and 22 older adults (age range: 65–75 years).

**Study 2:** 25 young and 30 older adult participants were recruited in Study 2. Five young and six older participants were excluded from the initial sample due to not completing the scanning session because of claustrophobia or discomfort, technical difficulties, chance source memory performance, or incidental MRI findings. Additionally, two young and two older participants were excluded from the present analyses due to excessive movement during the T1-weighted scan. Four older adults participated in both studies; only their Study 1 data were included in the present analyses. Thus, the final sample for Study 2 included 18 young (age range: 18–30 years) and 18 older adults (age range: 65– 75 years).

### Experimental materials and procedure

Participants in both studies underwent fMRI during a memory test (Figure 1). In both cases, participants first encoded concrete noun-visual image pairs. The words were paired with images of scenes or faces in Study 1 and with scenes or scrambled images in Study 2. The participants were required to imagine a scenario in which the object depicted by the word interacted with its paired image. They then rated the vividness of the imagined scenario on a 3-point scale (‘not vivid’ – ‘somewhat vivid’ – ‘very vivid’). In the subsequent retrieval phase, studied and novel words were employed as test items in a two-step source memory task. The requirement was to indicate whether the word was new or old and, if endorsed old, to judge the image category the word had been paired with at study (i.e., a scene vs. a face in Study 1; a scene belonging to a designated ‘target’ category vs. a non-target scene or scrambled image in Study 2).

**Figure 1:**
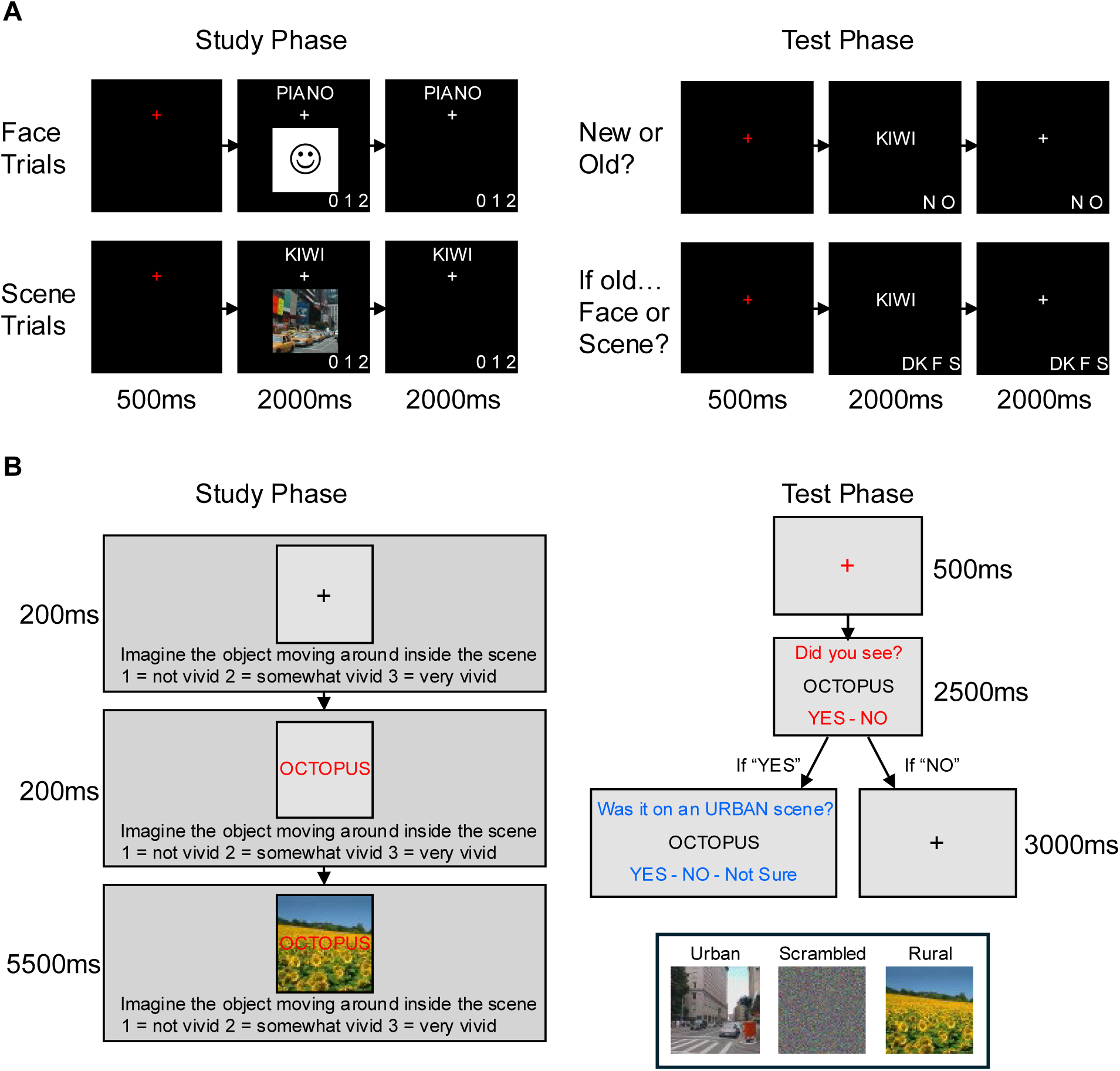
Overview of encoding and subsequent memory tasks for (A) Study 1 and (B) Study 2.

**Study 1:** Participants in Study 1 underwent fMRI during both the encoding and retrieval phases. Experimental items consisted of 288 concrete nouns paired with 96 images of faces and 96 images of scenes. The study phase was undertaken across four scanning runs, each containing 48 trials comprising words paired with an image (50% faces, 50% scenes) and 24 null trials, where a white fixation cross remained on the screen for the entire trial. A red fixation cross was presented for 500 ms at the commencement of each trial. The word-image pair remained on the screen for 2000 ms, followed by a white fixation cross for another 2000 ms.

The test phase was also undertaken across four scanning runs, each comprising 48 studied words, 24 new words, and 24 null trials. Each trial began with a red fixation cross presented for 500 ms, the test word for 2000 ms, and a white fixation cross for another 2000 ms. Participants were instructed to indicate whether the word was “Old” or “New” for each trial. If the endorsement was “Old,” participants were then required to make a source memory judgment, indicating whether the word had been paired with an image of a face or a scene. A “don’t know” option was included to discourage guessing.

**Study 2:** Participants in Study 2 underwent fMRI during the retrieval phase only. The out-of-scanner study task employed 240 concrete nouns, 120 color images of scenes (50% urban, 50% rural), and 60 scrambled (pixelated) scenes. Word-image pairs were presented in one of three randomized display locations: 20 on the left side of the screen, 20 in the middle, and 20 on the right. Each study phase trial began with a black fixation cross for 200 ms in the location corresponding with the upcoming word-image pair. The study word was then presented, joined by the background image 200 ms later. The word-image pair remained on the screen for 5500 ms.

The test phase was undertaken across five scanner runs, each divided into two mini-blocks. In the different mini-blocks, memory was tested for either the studied background or the studied location of the test word. The analyses described below focus only on scene reinstatement associated with background judgments. Each background task block contained 24 critical trials, comprising 18 studied word-image pairs and six new trials, intermixed with eight null trials. Each trial began with a red fixation cross presented for 500 ms, followed by the test word, which remained on the screen for 2500 ms.

During this time, participants indicated whether they recognized the word from the study phase by responding “Yes” or “No” to the prompt “Did you see?”. For test items endorsed as studied, participants were then required to respond “Yes,” “No,” or “Not Sure” to either the prompt “Was it on a rural scene?” or “Was it on an urban scene?” Task prompts were counterbalanced across participants such that any given participant received only one of the two prompts for the duration of the study.

#### MRI data acquisition and preprocessing

All structural and functional MRI data were acquired with the same Philips Achieva 3T MRI scanner (Philips Medical Systems, Andover, MA) equipped with a 32-channel head coil. Functional images were acquired using a T2*-weighted, blood oxygen level-dependent echoplanar imaging (EPI) sequence (sensitivity encoding [SENSE] factor 2, flip angle = 70 deg., TE = 30ms, FOV = 24 cm). EPI volumes from Study 1 consisted of 34 slices with a voxel size of 3 × 3 × 3 mm acquired in ascending order (TR = 2s, 1 mm interslice gap). In Study 2, EPI volumes comprised 44 slices acquired in an interleaved order at a voxel size of 2.5 × 2.5 × 2.5 mm (TR = 1.6s, 0.5 mm interslice gap) and a multiband factor of 2. T1-weighted anatomical images in both studies were acquired with a magnetization-prepared rapid gradient echo (MPRAGE) pulse sequence (FOV = 240 × 240, 1 × 1 × 1 mm isotropic voxels, 160 slices, sagittal acquisition). In the case of one young and eight older participants, the fMRI and MPRAGE data were acquired on separate days, with a median of 5.2 months between the two visits.

The fMRI data were preprocessed using Statistical Parametric Mapping (SPM12, Wellcome Department of Cognitive Neurology, London, UK) and custom MATLAB code. Functional images were realigned to the mean EPI image, and slice time was corrected using sinc interpolation to the middle slice. The images were then reoriented along the anterior and posterior commissures and spatially normalized to MNI space using a sample-specific EPI template following the procedures described by de Chastelaine et al. (2011, 2016). During the normalization step, the functional images from Study 2 were downsampled to a voxel size of 3 × 3 × 3 mm, thus matching the resolution of the functional images acquired in Study 1.

### Structural MRI analysis

#### Estimation of whole-brain cortical thickness

Estimates of whole-brain cortical thickness were derived from the T1-weighted image of each participant using Freesurfer’s (v6.0) semi-automatic processing pipeline (https://surfer.nmr.mgh.harvard.edu/fswiki; Dale et al. 1999; Fischl and Dale 2000; Fischl et al. 2002). Each 3-dimensional T1 volume underwent skull-stripping and intensity normalization, followed by reconstruction as an inflated 2-dimensional surface to identify white matter and pial surfaces. White matter and pial surfaces were visually inspected by one of six trained raters. All raters were blind to participant age. The raters corrected tissue identification errors by editing the brain masks returned by the first pass of the automated analysis. ‘Control points’ were also added to voxels on white matter paths that the Freesurfer pipeline had not appropriately labeled. Brain regions frequently requiring manual edits included the boundaries between grey matter and the pial surface in the temporal cortex, insula, and orbitofrontal cortex, and the delineation between the cortex and the cerebellum. A second rater inspected each scan for quality control purposes. Cortical thickness was defined as the distance between the pial surface and the grey/white matter boundary on a vertex-by-vertex basis for the cortical mantle of each hemisphere (de Chastelaine et al. 2023). Each hemisphere’s mean cortical thickness values were averaged to obtain mean whole-brain cortical thickness.

#### Estimation of whole-brain cortical volume and total intracranial volume

The “CortexVol” and “estimated Total Intracranial Volume” measures output by Freesurfer were used as estimates of whole-brain cortical volume and total intracranial volume (ICV), respectively. Before statistical analysis, whole-brain cortical volume estimates were adjusted for total intracranial volume to account for individual differences in head size. A linear regression was performed to make this adjustment, with whole-brain cortical volume as the dependent variable and total intracranial volume as the independent variable. Residuals were computed by subtracting the predicted cortical volume (based on total ICV) from the actual cortical volume for each participant. These adjusted cortical volume estimates were used as the primary volume variable for subsequent analyses. Hence, all references below to cortical volume refer to these adjusted estimates.

Note that we focus on whole-brain measures of cortical thickness and volume because of the evidence that regional structural measures tend to be intercorrelated and, individually, demonstrate unreliable associations with cognitive metrics (Salthouse et al. 2015; Kranz et al. 2018; Masouleh et al. 2019; Hou et al. 2021; Krogsrud et al. 2021; Masouleh et al. 2022; but, see Lee et al. 2016). Analyses that used structural metrics of cortical parcels that roughly correspond with the regions occupied by our regions of interest (ROIs) are reported in detail in the supplementary materials. At the request of a reviewer, we also examined the possibility that any associations that were identified between scene reinstatement and whole-brain structural metrics might be driven by structural characteristics of the prefrontal cortex. To this end, we included an additional three cortical parcels corresponding to the superior, middle, and inferior frontal gyri. Analyses involving these regions, including intercorrelations between the regional structural metrics and their associations with whole-brain cortical thickness and volume, are also detailed in the supplementary materials.

### Functional MRI analysis

#### ROI selection

We selected three scene-selective ROIs for the analyses, namely, the parahippocampal place area (PPA), the medial place area (MPA), and the occipital place area (OPA), as illustrated in Figure 2. The ROIs were derived from an unpublished independent localizer dataset in which participants viewed images of objects, faces, and scenes. The ROIs were defined using an across-participant scene > object + face contrast at a height threshold of p < 0.05 FWE-corrected. ROI definition followed that of Srokova et al. (2022). The PPA was defined by restricting the scene-selective cluster to the fusiform and parahippocampal gyrus labels in the Neuromorphometrics atlas available in SPM12.

**Figure 2:**
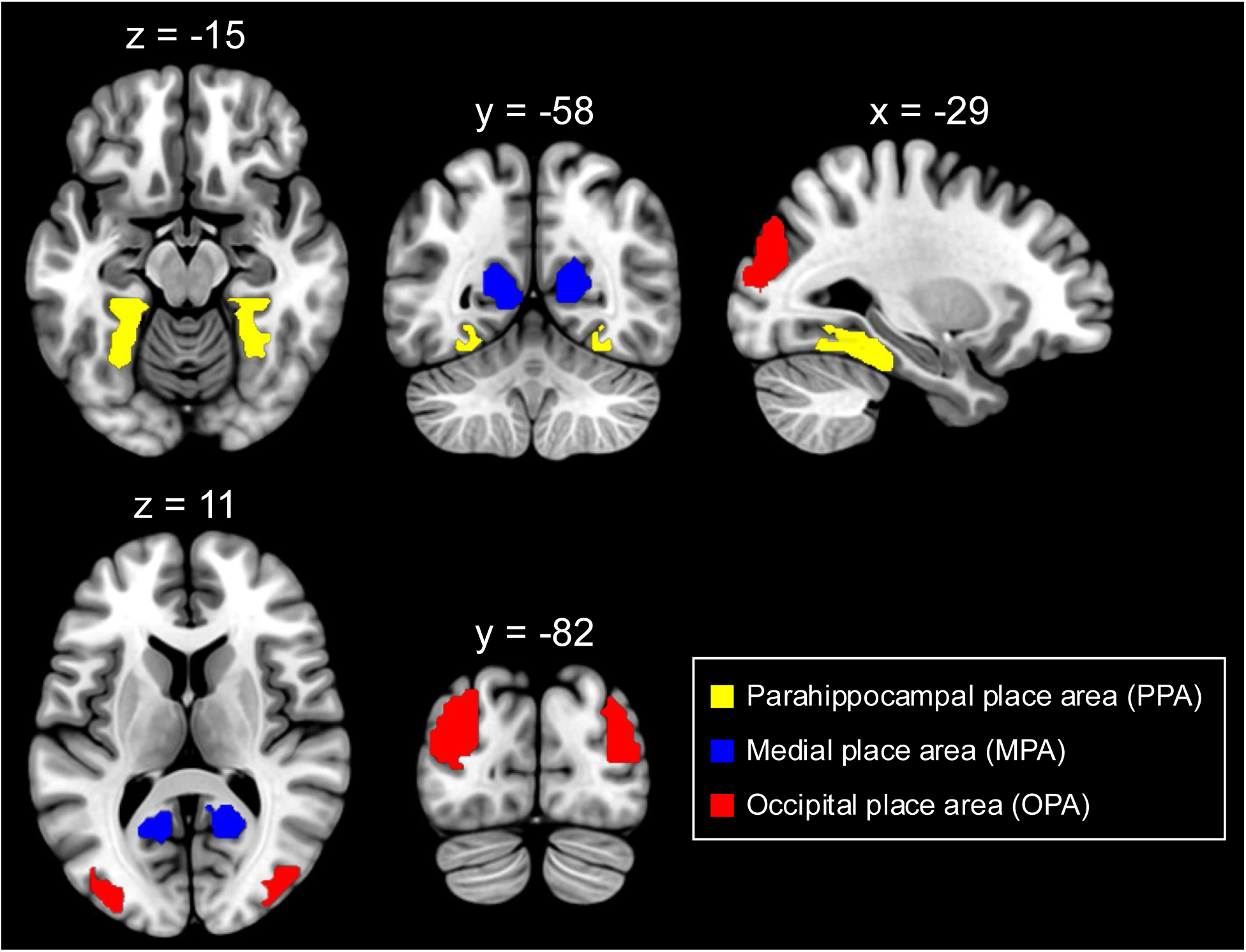
Regions of interest. Reinstatement was estimated in the parahippocampal place area (PPA), medial place area (MPA), and occipital place area (OPA). The regions are overlaid on the Montreal Neurological Institute (MNI) brain using MRIcroGL (https://www.nitrc.org/projects/mricrogl; Brett et al. 2001), with the PPA indicated in yellow, the MPA in blue, and the OPA in red. Slices are displayed in neurological convention at the following coordinates: axial slices at z = -15 and 11, coronal at y = -58 and -82, and sagittal at x = -29.

The OPA was defined by restricting the scene-selective cluster to the inferior and middle occipital gyrus Neuromorphometrics labels. As Neuromorphometrics does not provide an accurate anatomical mask for the MPA, that ROI was defined by inclusively masking the contrast with the outcome of a search in the Neurosynth database using the term “retrosplenial” (search in August 2019; search results were FDR-corrected at *p* < 0.00001; Yarkoni et al. 2011). The number of voxels contained within each ROI was 138 and 130 (left and right PPA, respectively), 117 and 135 (left and right MPA), and 238 and 204 (left and right OPA).

### Reinstatement index

The unsmoothed fMRI data from each participant were concatenated and subjected to a least-squares-all GLM (Rissman et al. 2004; Mumford et al. 2014) in which each trial is modeled with its own regressor in a single design matrix. Every trial was modeled with a delta function time-locked to the onset of the test item and convolved with SPM’s canonical hemodynamic response function. Covariates of no interest comprised six motion regressors reflecting rigid body translation and rotation, and regressors modeling the session means.

The single-trial blood-oxygen-level-dependent (BOLD) parameter estimates obtained from the GLM were employed to compute participant-specific reinstatement indices that quantified the strength of scene reinstatement in each ROI. We extracted the average across-voxel BOLD amplitude for each trial in each ROI, collapsing across the hemispheres. The reinstatement index was computed by subtracting the across-trial mean BOLD responses elicited by test words paired with scrambled images or faces from the mean response elicited by test words paired with the scenes, divided by their pooled standard deviation. Only trials associated with correct source judgments were employed to estimate these indices.

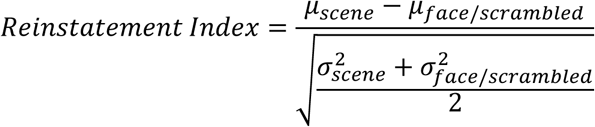

A positive reinstatement index signifies a greater mean BOLD response for words paired with scenes than for words paired with the alternate image category. Thus, the reinstatement index quantifies retrieval-related neural reinstatement of scene-related information. Of importance, because of the scaling function, the reinstatement index is insensitive to between-participant and between-group differences in the gain of the hemodynamic response function (see, for example, Liu et al. 2013). We elected to employ this univariate metric rather than a multi-voxel metric, such as pattern similarity, because of prior findings that the metric gave rise to larger effect sizes for reinstatement effects than did a pattern similarity metric and was more robustly associated across participants with memory performance (Srokova et al. 2021).

### Statistical Analysis

Statistical analyses were conducted using SPSS v. 30.0.0.0. In any case where a data point deviated by > 2.5 standard deviations from the sample mean, the analyses were re-run after excluding the relevant participant. Results are reported for the entire dataset unless the removal of the outlier changed the outcome of the analysis.

To examine age group differences in neural reinstatement, a mixed-effects ANOVA was performed with age group (young, older) as a between-subjects factor and ROI (PPA, MPA, OPA) and study (Study 1, Study 2) as within-subject factors. To examine the age differences in whole brain cortical volume and thickness, separate one-way ANOVAs were conducted with age group (young, older) as a between-subjects factor.

The relationships between age, neural reinstatement, whole-brain cortical thickness, and whole-brain cortical volume (adjusted for total intracranial volume) were examined using a series of multiple regression analyses employing the factors of thickness, volume, age group, study, and their interactions as predictors of reinstatement. For the analyses predicting reinstatement, the general form of the regression models was as follows:

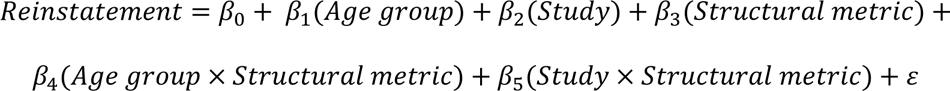

When behavioral performance was the dependent variable, the models were in the form:

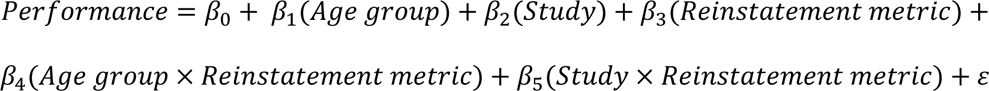

In any model where the interaction term involving either the age or study factor was non-significant, the term was dropped and the model was re-run.

## Results

### Behavioral results

The behavioral results from the two studies have been reported previously (Hill et al. 2021; Srokova et al. 2021) and are only briefly summarized below. Item recognition (*Pr*) was defined as the difference between the probability of a recognition hit and a false alarm. In Study 1, item recognition was lower in older than in young adults (p = .003), whereas no age differences were identified in Study 2 (p < .218; see Table 2).

**Table 2:**
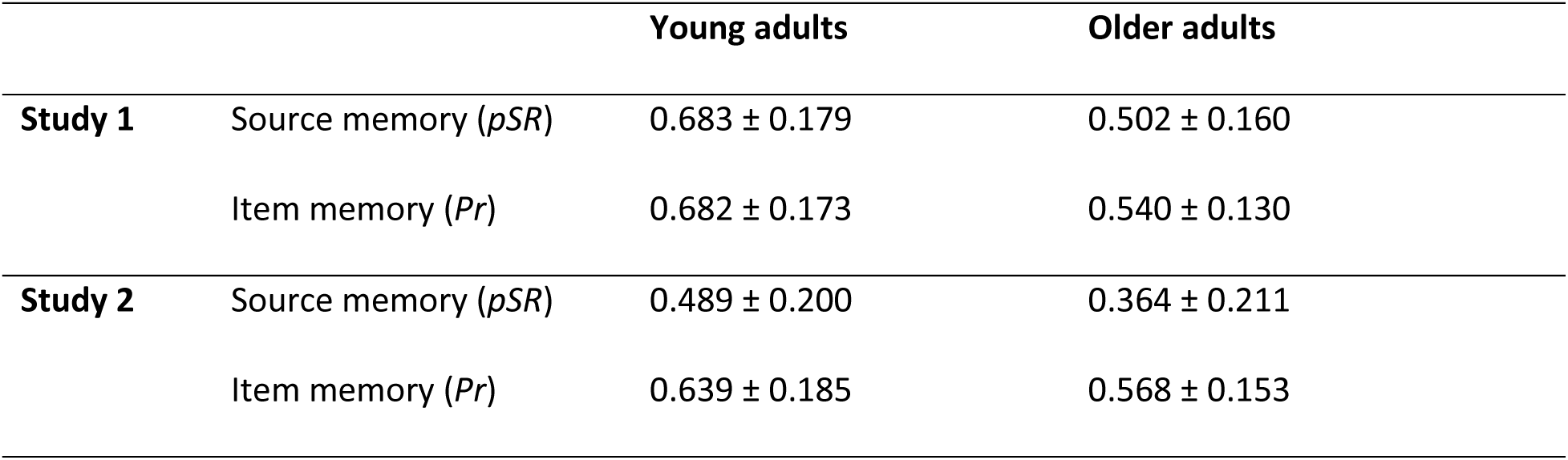
Item and source memory (M ± SD)

Source memory performance, operationalized as the probability of source recollection (*pSR*), was defined according to a single high-threshold model (Snodgrass and Corwin 1988) that included a correction for guessing (Park et al. 2008):

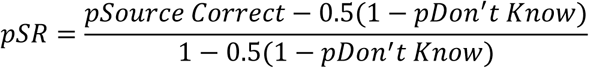

“*pSource Correct*” and “*pDon’t Know*” refer to the proportion of correctly recognized old trials associated with correct source judgments or a don’t know response, respectively. *pSR* was significantly lower in older than in young adults in the Study 1 sample (p < .001), but the age group differences in Study 2 did not reach significance (p = .075; see Table 2).

### Age differences in scene reinstatement

A 2 (age group) × 3 (ROI) × 2 (study) mixed effects ANOVA was conducted on the reinstatement indices derived from the PPA, MPA, and OPA ROIs, the outcome of which is reported in Table 3 and illustrated in Figure 3. The ANOVA revealed significant main effects of age group and ROI, while the study main effect and all 2-way and 3-way interactions failed to reach significance. The main effect of ROI reflected weaker reinstatement in the OPA than in either the PPA or the MPA (p < .001 in both cases). The main effect of age indicated that reinstatement was robustly weaker in the older adults than in the young adults in all ROIs and in both studies. Reinstatement effects were reliably greater than zero in all ROIs in the young participants (all p < .001), and in the PPA and MPA, but not the OPA, in older participants (ps = .011, .001, and .368, respectively).

**Figure 3:**
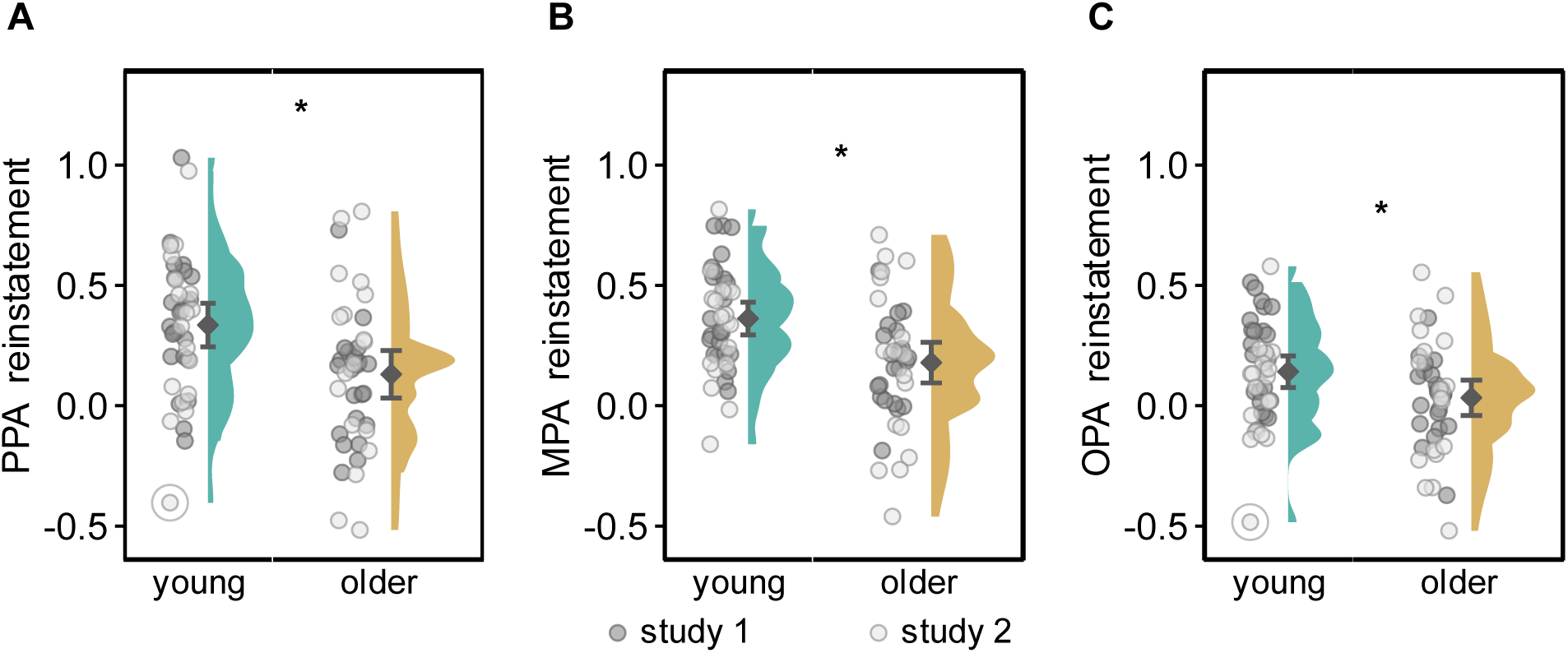
Distribution of reinstatement indices in Study 1 and Study 2 by age group in (A) the PPA, (B) the MPA, and (C) the OPA. Black dots and bars represent means and ±95% confidence intervals. Significant group differences are indicated by asterisks. Removal of outliers (circled) did not change the outcome of the statistical analyses.

**Table 3:** Outcome of study × age group × ROI ANOVA on reinstatement.

| | **F (1,78)** | $\eta_p^2$ | **p-value** |
| --- | --- | --- | --- |
| Study | 0.429 | .005 | .515 |
| Age group | 11.870 | .132 | **<.001\*** |
| ROI | 22.341 | .223 | **<.001\*** |
| Study × age group | 1.564 | .020 | .215 |
| Study × ROI | 1.881 | .024 | .174 |
| Age group × ROI | 2.318 | .029 | .132 |
| Study × age group × ROI | 0.026 | .000 | .872 |

Note: All F-tests had degrees of freedom of (1, 78). $\eta_p^2$ = partial eta squared. Significant effects ( $p < .05$ ) are shown in bold.

### Age differences in cortical thickness and volume

We employed 2 (study) × 2 (age group) between-subjects ANOVAs to examine age group differences in cortical thickness and volume estimates. The outcomes of these analyses are reported in Table 4 and illustrated in Figure 4.

**Figure 4:**
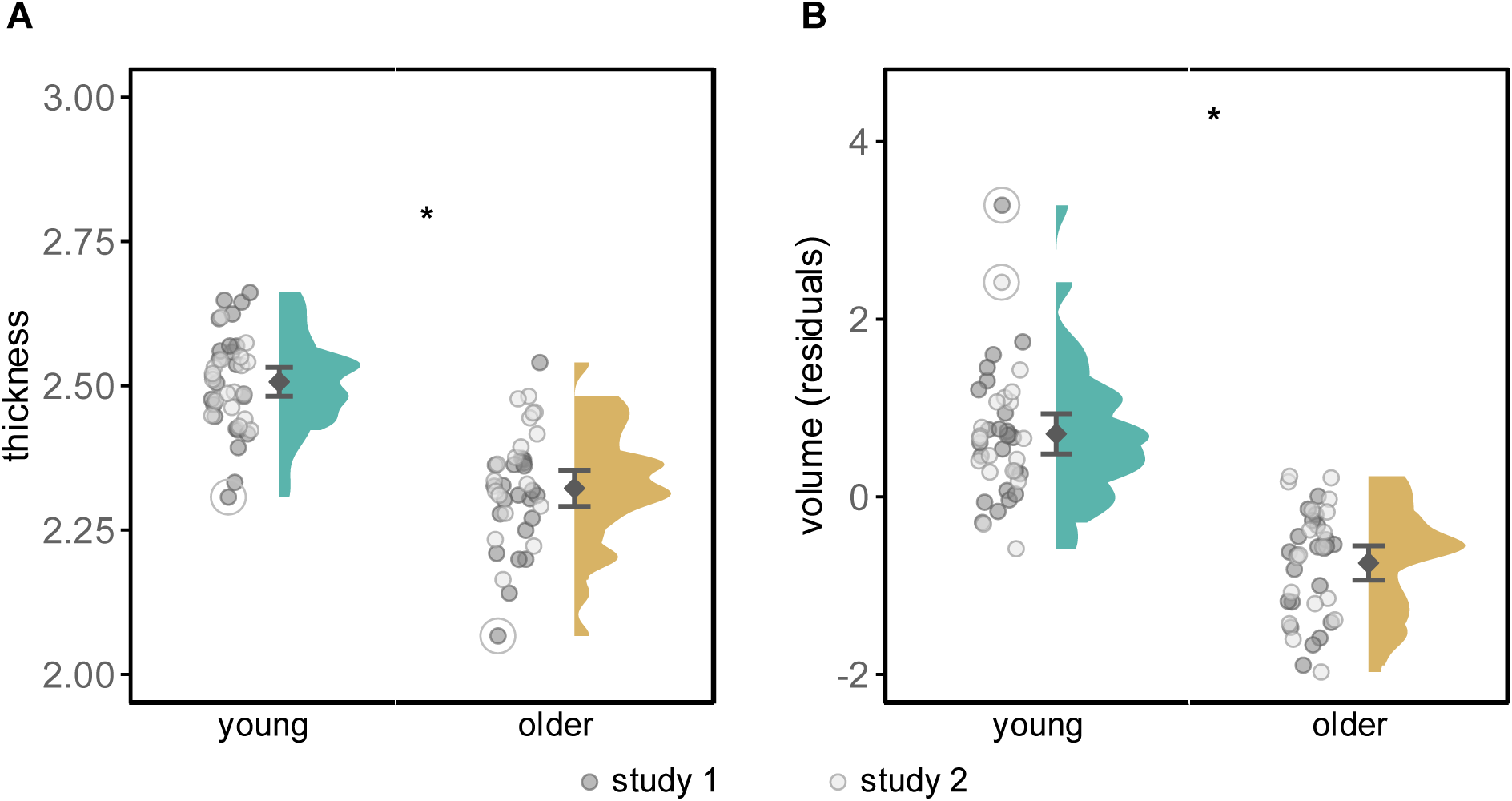
Distribution of estimates by age group of (A) cortical thickness and (B) cortical volume, combined across Study 1 and Study 2. The volumetric data are residualized for both ICV and study. Black dots and bars represent means and ±95% confidence intervals. Significant group differences are indicated by asterisks. Removal of outliers (circled) did not change the outcome of the statistical analyses.

**Table 4:** Outcome of the study × age group ANOVAs of cortical thickness and volume.

|  |  | **F (1 ,81)** | **$\eta_p^2$** | **p-value** |
| --- | --- | --- | --- | --- |
| **Cortical thickness** | Study | 1.595 | .020 | .210 |
|  | Age group | 84.666 | .520 | **<.001\*** |
| | Study × age group | 2.288 | .028 | .134 |
| **Cortical volume** | Study | 8.679 | .100 | **.004\*** |
|  | Age group | 94.128 | .547 | **<.001\*** |
| | Study × age group | 0.412 | .005 | .523 |

Note: All F-tests had degrees of freedom of (1, 81). $\eta_p^2$ = partial eta squared. Significant effects ( $p < .05$ ) are in bold.

As evident from Table 4, the ANOVA of thickness estimates revealed a significant main effect of age group, whereby thickness was lower in the older than the young adult group. Neither the main effect of study nor the age group × study interaction was significant. Thus, neither the thickness estimates nor the moderating effect of age differed significantly between the two studies. Paralleling the thickness analysis, the ANOVA of volume estimates revealed a significant main effect of age group, such that volume was lower in the older than the young adult group. There was also a significant main effect of study, with mean volume estimates lower in Study 1 than in Study 2; the reason for this difference is unclear. The age group × study interaction was not significant.

### Cortical thickness as a predictor of scene reinstatement

Multiple regression was employed to examine the relationships between age group, cortical thickness, and scene reinstatement in each ROI. As detailed in the methods, the initial regression models employed age group, study, thickness, the interactions between age group and thickness, and study and thickness as predictors. The interaction terms were non-significant in all cases (PPA model: age × thickness: p = .391, study × thickness: p = .998; MPA model: age × thickness: p = .939, study × thickness: p = .265; OPA model: age × thickness: p = .999, study × thickness: p = .318), and hence each model was re-run after excluding them. The outcomes from these reduced models are presented in Table 5, where it can be seen that thickness was positively correlated with reinstatement in the PPA only (see Figure 5).

**Figure 5:**
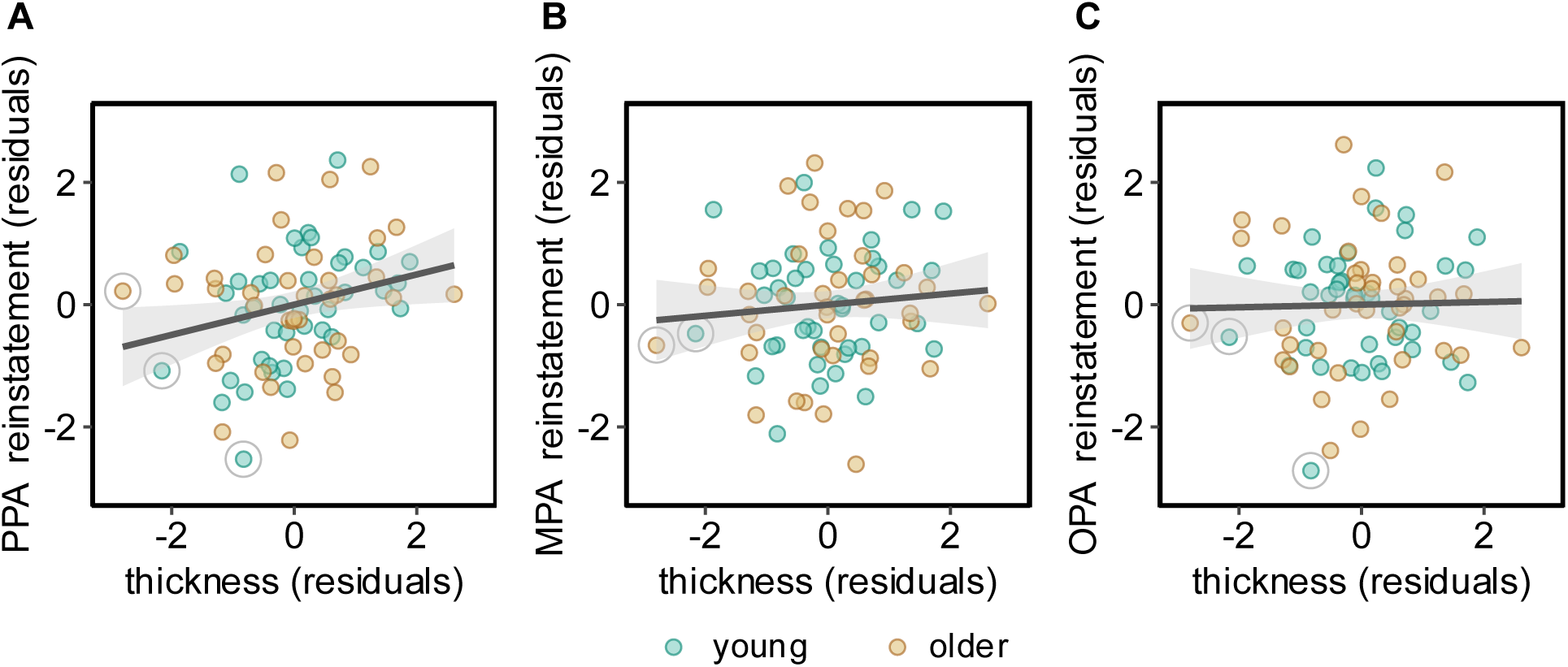
Scatter plots of the relationships between cortical thickness and scene reinstatement in the **(A)** PPA, **(B)** MPA, and **(C)** OPA. The shaded area around each regression line represents the 95% confidence interval. Removal of outliers (circled) did not change the outcome of the statistical analyses.

**Table 5:** Outcome of linear regression analyses testing whether cortical thickness, age group, and study significantly predicted reinstatement in the PPA, MPA, and OPA.

| | | *b* | SE *b* | $\beta$ | t | Partial corr. | p-value |
| --- | --- | --- | --- | --- | --- | --- | --- |
| **PPA reinstatement** | Cortical thickness | .836 | .372 | .340 | 2.251 | .247 | **.027\*** |
|  | Age group | -.051 | .095 | -.081 | -.535 | -.060 | .594 |
|  | Study | -.004 | .066 | -.007 | -.065 | -.007 | .949 |
| **MPA reinstatement** | Cortical thickness | .248 | .307 | .124 | .808 | .091 | .422 |
|  | Age group | -.136 | .078 | -.267 | -1.748 | -.194 | .084 |
|  | Study | -.042 | .055 | -.081 | -.761 | -.086 | .449 |
| **OPA reinstatement** | Cortical thickness | .053 | .278 | .030 | .192 | .022 | .848 |
|  | Age group | -.097 | .071 | -.216 | -1.370 | -.153 | .175 |
|  | Study | -.075 | .050 | -.166 | -1.516 | -.169 | .134 |

Note: *b* = unstandardized coefficient; SE *b* = standard error of the unstandardized coefficient; $\beta$ = standardized coefficient. Significant effects ( $p < .05$ ) are in bold.

### Cortical volume as a predictor of scene reinstatement

We used an analogous regression approach to examine the relationships between cortical volume and reinstatement in the PPA, MPA, and OPA. The interaction terms were again non-significant in all three cases (PPA model: age × volume: p = 0.941, study × volume: p = 0.493; MPA model: age × volume: p = 0.353, study × volume: p = 0.845; OPA model: age × volume: p =.655, study × volume: .251) and hence were dropped from the final models. The outcomes of these models are described in Table 6 and illustrated in Figure 6. As is evident from the table, cortical volume was a significant predictor of reinstatement in the PPA and MPA, but not the OPA.

**Figure 6:**
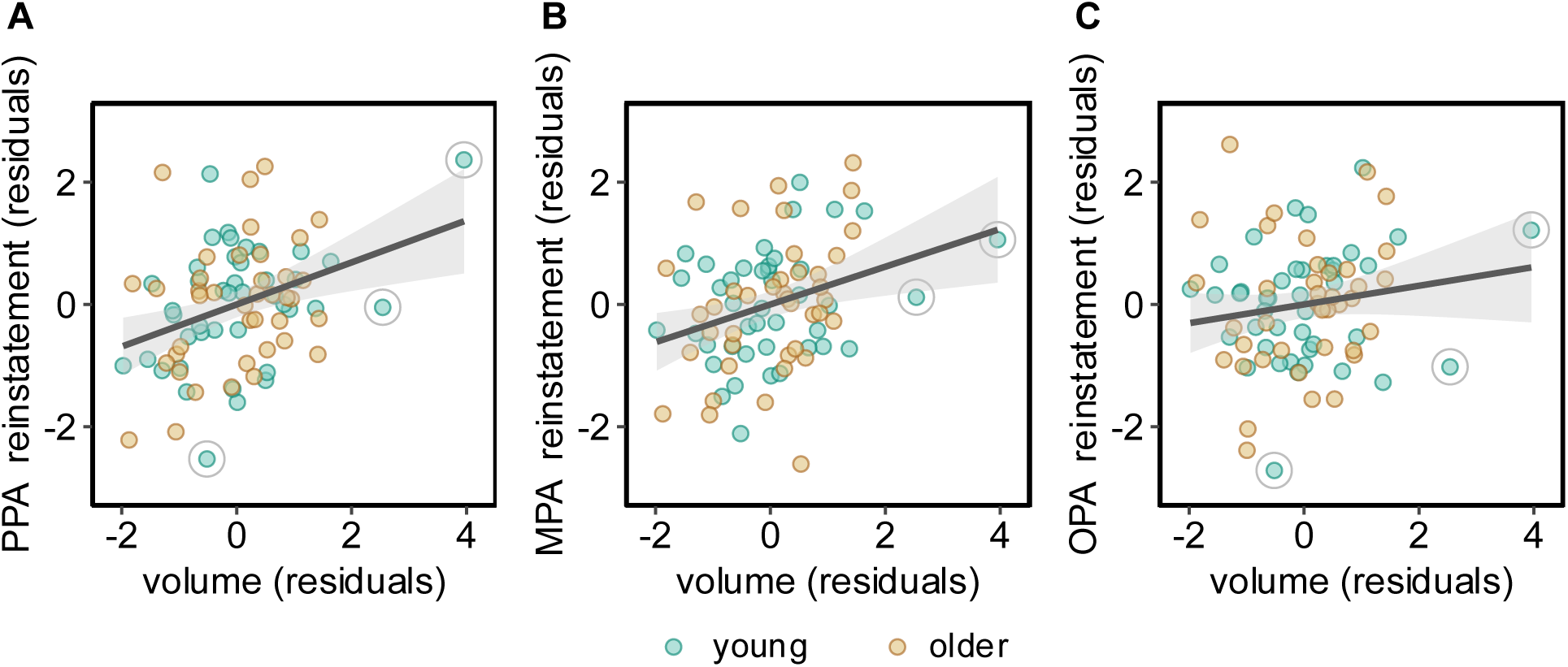
Scatter plots of the relationships between cortical volume and scene reinstatement in the **(A)** PPA, **(B)** MPA, and **(C)** OPA. The shaded area around each regression line represents the 95% confidence interval. Removal of outliers (circled) did not change the outcome of the statistical analyses.

**Table 6:** Outcome of multiple linear regression analyses testing whether cortical volume, age, and study significantly predicted reinstatement in the PPA, MPA, and OPA.

| | | *b* | SE *b* | $\beta$ | *t* | Partial corr. | p-value |
| --- | --- | --- | --- | --- | --- | --- | --- |
| **PPA reinstatement** | Cortical volume | .158 | .049 | .497 | 3.238 | .344 | **.002\*** |
|  | Age group | .022 | .094 | .035 | .233 | .026 | .816 |

CORTICAL STRUCTURE AND AGE DIFFERENCES IN NEURAL REINSTATEMENT
|  | Study | -.052 | .067 | -.082 | -.779 | -.088 | .439 |
| --- | --- | --- | --- | --- | --- | --- | --- |
| **MPA reinstatement** | Cortical volume | .113 | .040 | .439 | 2.865 | .309 | **.005\*** |
|  | Age group | -.019 | .077 | -.037 | -.244 | -.028 | .808 |
|  | Study | -.084 | .054 | -.164 | -1.554 | -.173 | .124 |
| **OPA reinstatement** | Cortical volume | .051 | .037 | .224 | 1.368 | .153 | .175 |
|  | Age group | -.034 | .072 | -.211 | -.466 | -.053 | .642 |
|  | Study | -.096 | .051 | -.075 | -1.874 | -.208 | .065 |

Note: $b$ = unstandardized coefficient; SE $b$ = standard error of the unstandardized coefficient; $\beta$ = standardized coefficient. Significant effects ( $p < .05$ ) are in bold.

In a follow-up analysis, we examined whether cortical thickness and volume were independent predictors of reinstatement in the PPA, the only ROI where each metric correlated reliably with reinstatement. The outcome of the model is presented in Table 7, where it is evident that only volume accounted for a significant fraction of the variance in reinstatement.

**Table 7:** Outcome of multiple regression analyses predicting reinstatement in the PPA given the predictors cortical volume, cortical thickness, age group, and study.

| | | *b* | SE *b* | $\beta$ | *t* | Partial corr. | *p*-value |
| --- | --- | --- | --- | --- | --- | --- | --- |
| **PPA reinstatement** | Cortical volume | .134 | .054 | .422 | 2.465 | .248 | **.016\*** |
|  | Cortical thickness | .399 | .401 | .162 | .994 | .100 | .323 |
|  | Age group | .061 | .102 | .098 | .600 | .060 | .550 |
|  | Study | -.051 | .067 | -.081 | -.769 | -.087 | .444 |

Note: *b* = unstandardized coefficient; SE *b* = standard error of the unstandardized coefficient; $\beta$ = standardized coefficient. Significant effects ( $p < .05$ ) are in bold.

### Contributions of broadening and attenuation to age and volume effects on reinstatement

To further elucidate the findings for scene reinstatement, we analyzed the magnitudes of the BOLD responses elicited by the test words studied with scenes vs. the unpreferred image category (i.e., faces in Study 1, scrambled scenes in Study 2). As OPA reinstatement was non-significant in the older age group, these analyses were confined to the PPA and MPA. The data were subjected to a separate 2 (study) × 2 (age group) × 2 (image category) mixed-effects ANOVA for each ROI. As is evident in Table 8, these ANOVAs gave rise to main effects of study, age group, and image category in the PPA, and to main effects of study and image category in the MPA. The effects of age group and image category were modified by an age group × image category interaction in both the PPA and MPA. Figure 7 depicts the data pooled across experiments and ROIs, illustrating that the interaction reflected an absence of age differences in the responses elicited by the test items paired with scenes (p = .383) and a robust difference in favor of the older group in the responses to items paired with the unpreferred image categories (t80 = -3.85, p < .001, d = -.848; for a breakdown of these age group differences by ROI, see Table 9). Therefore, these findings indicate that the age-related attenuation of scene reinstatement reported above was a consequence of an age-related enhancement of the BOLD responses elicited by test items paired with the unpreferred images.

**Figure 7:**
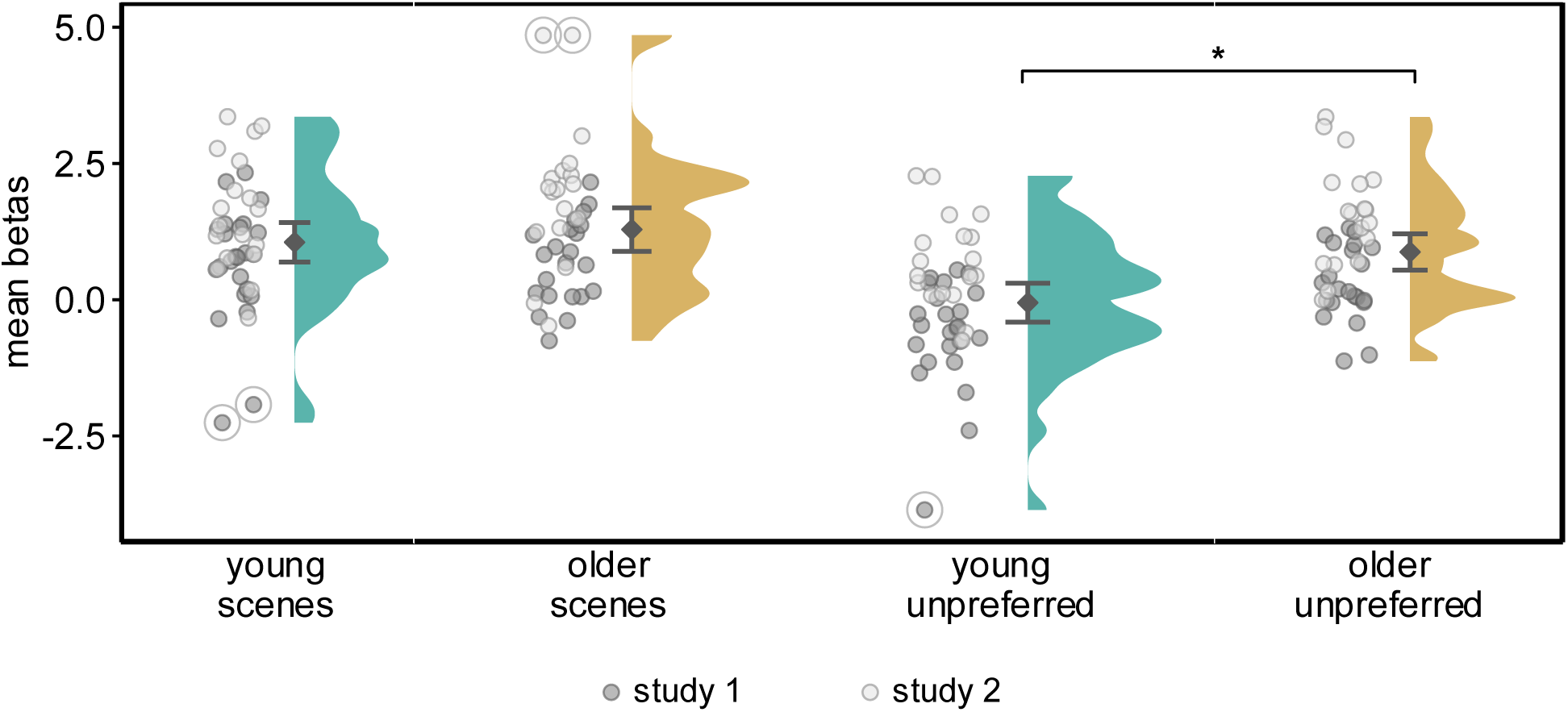
Plot of the mean beta estimates (collapsed across PPA and MPA ROIs) for test words studied with scenes and unpreferred image categories for the young and older age groups. Black dots and bars represent means and ±95% confidence intervals, and significant group differences are indicated by asterisks. Removal of outliers did not change the outcome of the statistical analyses.

**Table 8:** Outcome of study × age group × image category ANOVAs on BOLD response magnitudes at test for each ROI.

| | | **F (1, 78)** | $\eta_p^2$ | **p-value** |
| --- | --- | --- | --- | --- |
| **PPA** | Study | 23.192 | .229 | **<.001\*** |
|  | Age group | 11.066 | .124 | **.001\*** |
|  | Image category | 39.984 | .339 | **<.001\*** |
| | Study × age group | .012 | .000 | .913 |
| | Study × image category | .605 | .008 | .439 |
| | Age group × image category | 11.279 | .126 | **.001\*** |
| | Study × age group × image category | 1.851 | .023 | .178 |
| **MPA** | Study | 36.568 | .319 | **<.001\*** |
|  | Age group | 3.549 | .044 | .063 |
|  | Image category | 98.977 | .559 | **<.001\*** |
| | Study × age group | .364 | .005 | .548 |
| | Study × image category | 1.653 | .021 | .202 |
| | Age group × image category | 15.087 | .162 | **<.001\*** |
| | Study × age group × image category | .720 | .009 | .399 |

Note: All F-tests had degrees of freedom of (1, 78). $\eta_p^2$ = partial eta squared. Significant effects ( $p < .05$ ) are in bold.

**Table 9:** Independent samples t-tests showing age differences in BOLD responses elicited by the test words studied with scenes vs. the unpreferred image category, broken down by ROI.

|  |  | t | p-value | Cohen's d |
| --- | --- | --- | --- | --- |
| **PPA mean beta** | Scenes | -1.386 | .170 | -.306 |
|  | Unpreferred | .4.463 | **<.001\*** | -.979 |
| **MPA mean beta** | Scenes | -.382 | .704 | -.085 |
|  | Unpreferred | -2.806 | **.006\*** | -.621 |

Note: Significant effects ( $p < .05$ ) are in bold.

Using the same regression approach employed with the reinstatement index, we went on to examine the associations between the BOLD responses elicited by test items paired with scenes and the unpreferred image categories, age, study, and cortical volume, separately for the two ROIs. As previously, the age × functional metric interaction terms were dropped from the final models where they were non-significant. The results from each model are summarized in Table 10. In the case of the parameter estimates associated with scene trials, the variables of study, age group, and volume were all significant predictors of the size of the estimates.

**Table 10:** Outcome of linear regression analyses testing whether cortical volume, age group, and study significantly predicted BOLD activity elicited by scene trials or unpreferred image trials, broken down by ROI.

| ROI | Category | | *b* | SE *b* | $\beta$ | *t* | Partial corr. | p-value |
| --- | --- | --- | --- | --- | --- | --- | --- | --- |
| **PPA** | *Scenes* | Cortical volume | .442 | .174 | .385 | 2.541 | .277 | **.013\*** |
|  |  | Age group | .964 | .336 | .425 | 2.865 | .309 | **.005\*** |
|  |  | Study | .688 | .239 | .301 | 2.881 | .310 | **.005\*** |
|  | *Unpreferred* | Cortical volume | -.002 | .156 | -.001 | -.010 | -.001 | .992 |
|  |  | Age group | .993 | .302 | .433 | 3.292 | .349 | **.001\*** |
|  |  | Study | 1.031 | .214 | .447 | 4.821 | .479 | **<.001\*** |
| **MPA** | *Scenes* | Cortical volume | .579 | .207 | .402 | 2.792 | .301 | **.007\*** |
|  |  | Age group | .926 | .402 | .326 | 2.305 | .253 | **.024\*** |
|  |  | Study | 1.135 | .285 | .396 | 3.985 | .411 | **<.001\*** |
|  | *Unpreferred* | Cortical volume | .234 | .187 | .164 | 1.254 | .141 | .214 |
|  |  | Age group | 1.146 | .361 | .407 | 3.173 | .338 | **.002\*** |
|  |  | Study | 1.518 | .256 | .535 | 5.924 | .557 | **<.001\*** |

Note: *b* = unstandardized coefficient; SE *b* = standard error of the unstandardized coefficient; $\beta$ = standardized coefficient. Significant effects ( $p < .05$ ) are in bold.

Study and age group were significant predictors of the size of the responses elicited by the test items paired with the unpreferred categories in both ROIs, but cortical volume did not significantly predict these responses in either case. The outcomes of these analyses did not change with the removal of outliers. Thus, the relationship between scene reinstatement and cortical volume reported above reflects the mediating effect of volume on BOLD responses associated with scene retrieval.

### Cortical thickness, cortical volume, and reinstatement as predictors of memory performance

Lastly, we ran a series of multiple regressions to examine whether source memory performance (pSR) was predicted by reinstatement, cortical thickness, or cortical volume. pSR was residualized by study to remove study-specific differences in performance and hence provide a study-independent metric of memory performance. Age group and its interactions with study and reinstatement were included as predictors in the initial models. In no case were these interactions significant (ps > .340). As is evident from Table 11 (see also Figure 8), PPA reinstatement and age group were both significant predictors of memory performance. The association with reinstatement remained when cortical volume and thickness were added to the model. By contrast, neither MPA reinstatement nor OPA reinstatement significantly predicted memory performance (ps = .137, .413, respectively). Furthermore, after controlling for age group and study, neither cortical thickness nor volume were, by themselves, significant predictors of memory performance (r = .170 and .169, respectively, ps > .130).

**Figure 8:**
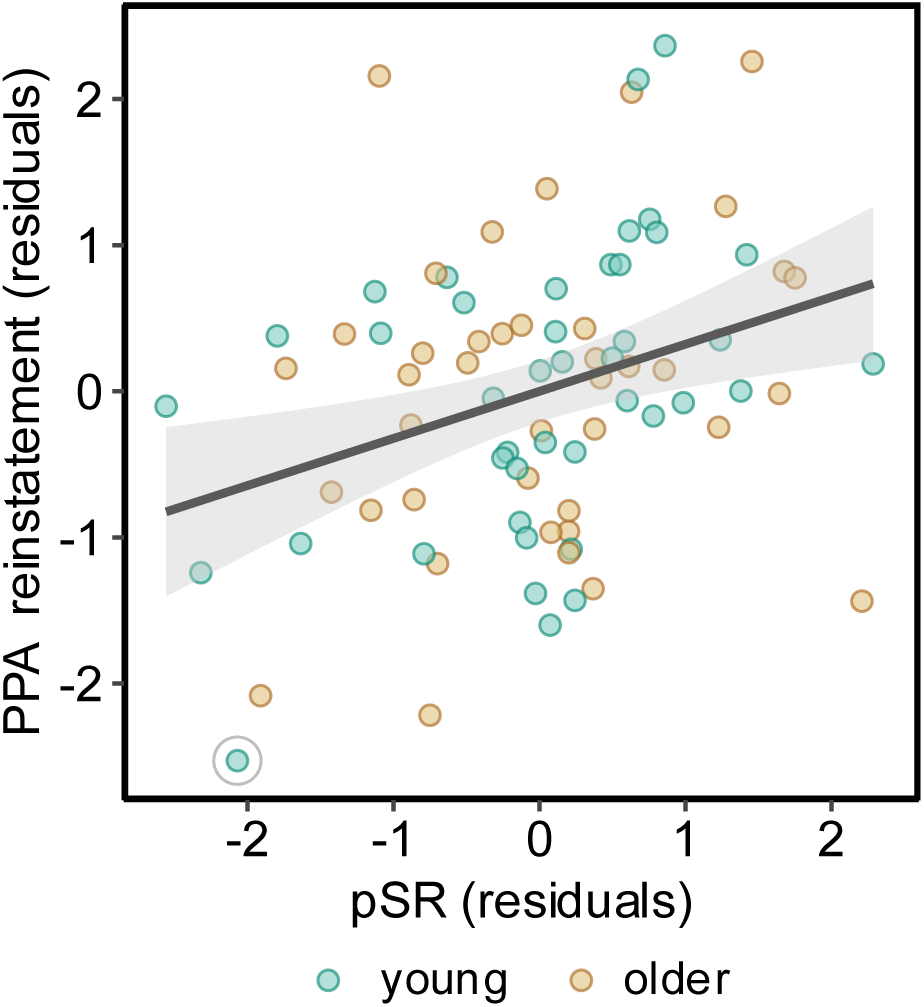
Scatter plot of the relationship between PPA reinstatement and memory performance (pSR) after controlling for age group and study. The shaded area around the regression line represents the 95% confidence interval. Removal of the outlier (circled) did not change the outcome of the statistical analyses.

**Table 11:** Outcome of multiple regression analyses predicting memory performance from the predictors of PPA reinstatement, age group, and study (upper) and PPA reinstatement, age group, study, cortical volume, and thickness (lower).

| | *b* | SE *b* | $\beta$ | *t* | Partial corr. | p-value |
| --- | --- | --- | --- | --- | --- | --- |
| PPA reinstatement | .968 | .328 | .307 | 2.949 | .317 | **.004\*** |
| Age group | -.593 | .206 | -.300 | -2.879 | -.310 | **.005\*** |
| Study | .002 | .196 | .001 | .009 | .001 | .993 |
| PPA reinstatement | .869 | .354 | .276 | 2.451 | .271 | **.017\*** |
| Cortical volume | .047 | .175 | .047 | .266 | .030 | .791 |
| Cortical thickness | .879 | 1.256 | .114 | .700 | .080 | .486 |
| Age group | -.383 | .319 | -.194 | -1.203 | -.137 | .233 |
| Study | -.038 | .209 | -.019 | -.183 | -.021 | .855 |

Note: *b* = unstandardized coefficient; SE *b* = standard error of the unstandardized coefficient; $\beta$ = standardized coefficient. Significant effects ( $p < .05$ ) are in bold.

## Discussion

We examined associations between retrieval-related neural reinstatement of scene information in the three scene-selective ROIs (PPA, MPA, and OPA), whole-brain cortical thickness and volume, and memory performance in samples of young and older adults. As expected, given prior findings (Rugg and Srokova 2024), there were robust age differences in reinstatement strength in each ROI. Also consistent with the prior literature, cortical volume and thickness were markedly lower in the older age group.

Multiple regression revealed associations between each of the structural metrics and scene reinstatement in the PPA and MPA, although the association with thickness was restricted to the PPA. Moreover, only volume accounted for a significant fraction of the variance in PPA reinstatement when the two structural metrics were included in the same regression model. Additionally, the effect of age on scene reinstatement in the PPA and MPA was fully mediated by cortical volume. Lastly, reinstatement strength in the PPA was associated with source memory performance independently of age and cortical volume. Below, we discuss the implications of these findings.

Before discussing the findings for the PPA and MPA, we address why, unlike in these ROIs, there were no detectable associations between OPA reinstatement and either whole brain or regional cortical thickness or volume. One possibility is that the mechanisms responsible for scene reinstatement in this region differ from those in the PPA and MPA, such that they are less sensitive to the factors captured by whole brain cortical volume that are responsible for modulating reinstatement in these regions (see discussion below). A more mundane possibility, however, is that OPA reinstatement metrics were too noisy and exhibited insufficient across-participant variability for a reliable association with volume to be detected at the statistical power afforded by the present study. In support of this possibility, we note that OPA reinstatement effects were markedly weaker than those in the PPA and MPA, to the extent that they did not even approach statistical significance in the older sample. A more highly powered study may help to arbitrate between these alternate accounts.

As noted above, the finding that older adults demonstrated weaker scene reinstatement than young adults is unsurprising, given prior reports. The present findings do, however, shed additional light on those prior findings, most prominently through the association that was identified between cortical volume and scene reinstatement in the PPA and MPA. These associations were robust and, strikingly, cortical volume fully mediated the association between age group and reinstatement in these ROIs (see Figure 9). Of importance, the relationship between scene reinstatement and cortical volume remained when the analysis was restricted to the young age group only (collapsing across the PPA and MPA, partial r = .377, p = .017, after controlling for chronological age and study; note that volume and chronological age also correlated reliably in this age group, partial r = -.327, p = .037).

**Figure 9:**
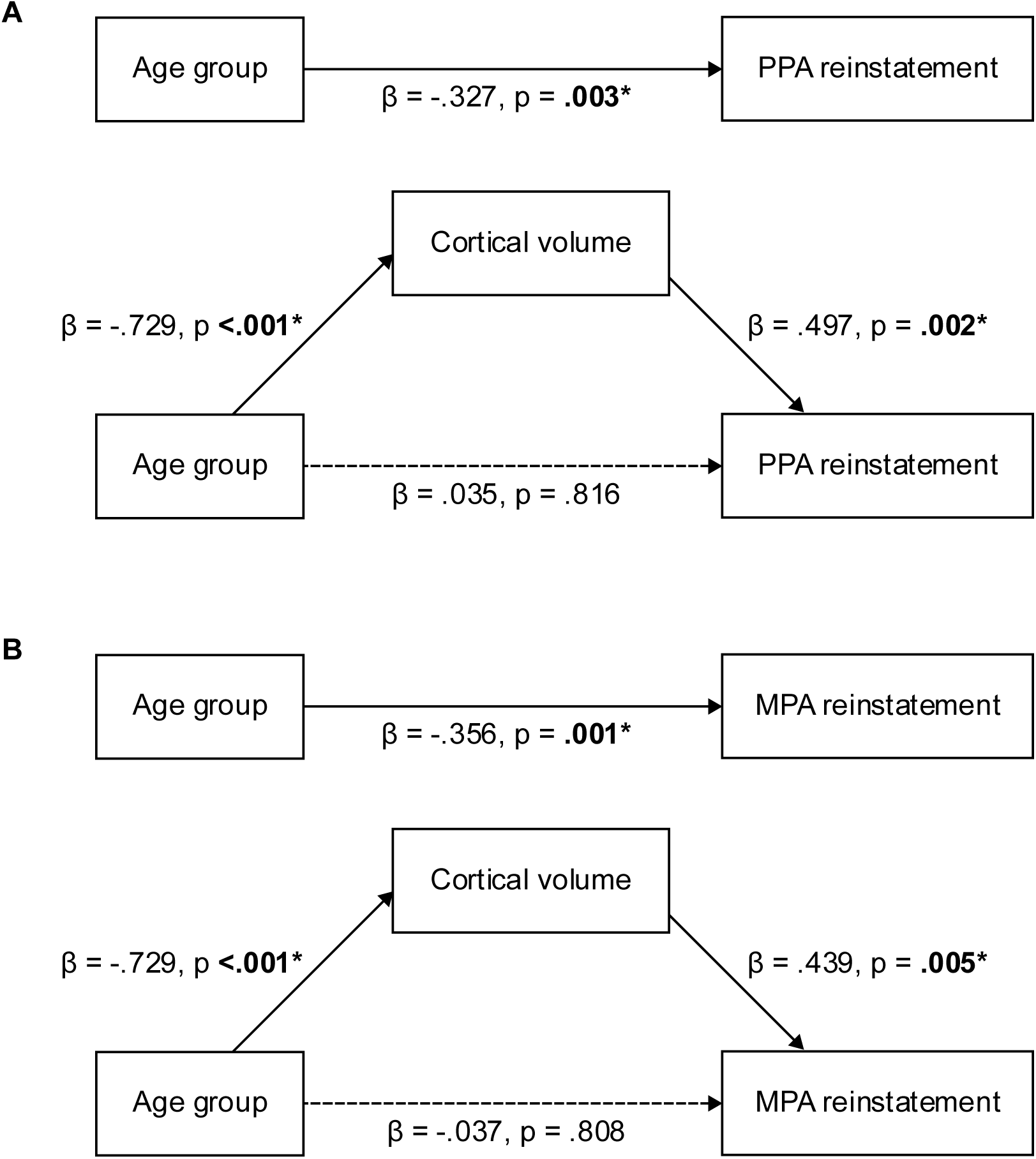
Path diagrams illustrating the mediating effect of cortical volume on the association between age group and scene reinstatement in the (A) PPA and (B) MPA. Standardized regression coefficients (β) and p-values are shown for each path. In both cases, the indirect effect was statistically significant as determined by bootstrapping with 5000 samples using the PROCESS macro (version 5.0) in SPSS (A: 95% CI = [-.3673, -.0644]; B: 95% CI = [-.2770, -.0544]). Study was included as a covariate of no interest in each model.

How does the finding that PPA and MPA scene reinstatement covaried not with age but with cortical volume contribute to the understanding of the determinants of reinstatement strength? One obvious implication is that prior findings of age-related reductions in scene reinstatement in these regions (e.g., Johnson et al. 2015; Trelle et al. 2020) may have arisen because of the confounding effects of cortical volume. This leads directly to the question: why should cortical volume be associated with reinstatement? A full answer to this question will require a more comprehensive understanding of the neurobiological determinants of MRI volumetric measures than exists currently. For example, elucidation of the relative contributions of different microstructural factors to cortical volumetric measures (such as neuron density, size and arborization, glial and myelin content, etc.) may bring us closer to an answer. The present findings do, however, offer some suggestions.

First, the finding that the association between reinstatement and cortical volume is age-invariant indicates that while the impact of age on reinstatement is likely attributable to the well-documented finding that cortical volume (and, indeed, total gray matter) declines monotonically across most of the human lifespan (Bethlehem et al. 2022), there is nothing in the present data to suggest that the association strengthens in later life. Thus, the factors responsible for the association are unlikely to include those underpinning the reductions in cortical volume caused by age-related pathologies like Alzheimer’s Disease or cerebrovascular disease (e.g., neuronal loss or white matter hyperintensities). Of course, this is not to say that reinstatement strength would not be impacted by such pathologies — this is a question that would be well worth pursuing—but given the findings from the young age group described above, neuropathological factors seem unlikely to be the mechanism underpinning individual differences in reinstatement strength in cognitively healthy adults, at least up to an age of around 75 years.

Another clue to the factors responsible for the association between reinstatement and cortical volume comes from the analyses conducted on the BOLD responses elicited by the test items paired with the scene versus the unpreferred image categories. These analyses revealed that volume was specifically associated with the functional activity elicited during scene retrieval. A speculative explanation for the specificity of this association is that volume acts as a constraint on the dynamic range of the BOLD responses elicited in scene-selective neural populations. Whether this constraining effect—should it exist—acts at the neural or the hemodynamic level is unclear.

As just alluded to, the analysis of the BOLD responses associated with the scene and unpreferred image categories clearly demonstrated that age-related attenuation of scene reinstatement is a consequence of neural ‘broadening’ (heightened responses to exemplars of an ‘unpreferred’ stimulus category) rather than neural ‘attenuation’ (diminished responses to exemplars belonging to a ‘preferred’ category). Both patterns of findings have been reported previously in studies describing age-related reductions in the selectivity of BOLD responses to preferred stimulus categories in category-selective cortical regions (e.g., Chee et al. 2006; Park et al. 2012; Koen et al. 2019; Srokova et al. 2020; for review, see Koen and Rugg 2019). However, few studies have examined this issue when category information is retrieved from memory rather than directly perceived (for a recent example, see Hill et al. 2021). The present findings are consistent with evidence (reviewed in Koen and Rugg 2019) that age-related reductions in neural selectivity are a consequence of broader ‘tuning’ functions in category-selective cortex and suggest that the proposal is equally applicable to the neural selectivity of mnemonic representations.

For the reasons noted in the introduction, we elected to use whole-brain structural metrics when conducting our primary analyses examining possible relationships between brain structure and scene reinstatement. As already discussed, the analyses identified a robust association between whole-brain cortical volume and reinstatement in both the PPA and MPA. As reported in the supplementary materials, when these analyses were repeated using as metrics the volumes of cortical regions that roughly corresponded with the regions occupied by our ROIs, no associations with reinstatement were evident. Additionally, reminiscent of findings from studies examining associations between regional structural metrics and cognition (e.g., Cox et al. 2019; Masouleh et al. 2019), weak associations identified between prefrontal regional volumes and reinstatement were no longer evident after controlling for the volume of non-frontal cortex. In short, we could find no evidence for regionally specific associations between scene reinstatement and structural metrics. Analogous null findings for regional associations between functional and structural metrics in cognitively healthy participants have been reported previously (de Chastelaine et al. 2016; Hou et al. 2020; Kidwai et al. 2025; see Mooraj et al. 2025 for review).

These null findings are open to at least two interpretations. First, they might signify that reinstatement is sensitive to a generic (trans-regional) characteristic of the cortex that is represented most reliably by the whole-brain measure. By this argument, the microstructural properties of specific cortical regions are less important as determinants of reinstatement strength than a generic factor captured by whole-brain volume. Alternately, reinstatement might indeed depend upon specific regional characteristics, but these are sufficiently highly correlated across regions to lead to an association at the whole-brain level, while the single-region metrics are insufficiently reliable to yield comparable findings at the statistical power afforded by the present experiment. Resolution of this issue will require research that employs larger sample sizes and more fine-grained structural analyses than those available here.

Whereas whole brain cortical volume demonstrated robust associations with PPA and MPA scene reinstatement, this was not the case for cortical thickness. As is evident from Table 5, thickness failed to demonstrate any association with MPA reinstatement (this was also the case for the OPA, where volume also failed to demonstrate a reliable association). And while thickness demonstrated a relatively weak association with reinstatement in the PPA, this association did not survive when volume and thickness were included in the same regression model. Why thickness proved to be an inferior predictor of reinstatement compared with volume is presently unclear, but this result is unlikely to be a consequence of excessive collinearity between the two metrics; controlling for age and study, they correlated across participants at r = .442, and thus shared less than 20% of their variance. Moreover, if the two metrics were largely redundant, thickness would have been expected to co-vary with reinstatement in both the PPA and the MPA. Clearly, the two metrics reflect dissociable neocortical characteristics, with those reflected by volume being the most salient in the case of retrieval-related neural reinstatement (cf. de Chastelaine et al. 2023).

Echoing several prior reports that metrics of neural selectivity derived from the PPA are associated with memory performance (e.g., Koen et al. 2019; Srokova et al. 2020; Srokova et al. 2024; see also Trelle et al. 2020), we identified a robust, age- and study-invariant across-participants relationship between PPA scene reinstatement and source memory performance. These findings are consistent with the proposal that the PPA plays a key role in the encoding and retrieval of episodic information, perhaps because of its contribution to the processing of mnemonically relevant contextual information (Eichenbaum et al. 2007; Aminoff et al. 2013). Of importance, unlike in the case of age, this association was not mediated by cortical volume (the correlation between volume and memory performance was r = .158, p = .162, controlling for age group and study). Thus, the variance in scene reinstatement shared with memory performance was independent of the variance shared with cortical volume (and, by extension, age group). Identifying the source—or sources—of the first of these different variance components could help elucidate the determinants of individual differences in memory function.

### Limitations

The present study shares some of the limitations of the two studies that contributed to its dataset. One important limitation comes from the employment of cross-sectional designs. Consequently, any effects of age group cannot be attributed unambiguously to the effects of aging rather than to one or more confounding factors such as selection bias or cohort effects (Rugg 2017). A second limitation is the relatively small sample size. Whereas this is, of course, larger than those of the contributing studies, our analyses remain underpowered to detect small effects, such as those that might exist between regional structural metrics and scene reinstatement. Lastly, while the reinstatement metric that was employed in our primary analyses should be impervious to age and individual differences in the gain of the hemodynamic transfer function, this is not the case for the analyses of the BOLD responses that contributed to the metric. Therefore, we cannot rule out the possibility that some of the age differences identified in those analyses are attributable to age differences manifest at the neurovascular rather than the neural level.

### Conclusions

The present findings advance understanding of the impact of age on retrieval-related neural reinstatement of scene information in two major ways. First, the findings suggest that prior reports of age differences in scene reinstatement strength are likely due to the confounding influence of cortical volume. Here, we found that after controlling for volume, age explained only a trivial and non-significant fraction of the variance in reinstatement in two of the three ROIs that were examined. Second, the robust across-participants association we identified between PPA reinstatement strength and memory performance was unmodified by either cortical volume or age group. Therefore, we conclude that two independent factors contribute to across-participant variability in reinstatement strength. One of these factors is reflected in brain structural metrics such as cortical volume, which explains the association between reinstatement and age. The other factor, the determinants of which are unknown, is associated with some aspect of cognitive function that contributes to individual differences in performance on the experimental memory test (and, perhaps, performance on other cognitive tests also; cf. Trelle et al. 2020).

It will be of considerable interest to see how well the present findings generalize to other experimental settings and to memoranda that elicit neural reinstatement in other category-selective cortical regions.

## Acknowledgments

We gratefully acknowledge the contributions of Marianne de Chastelaine, Mingzhu Hou, Amber Kidwai, Seham Kafafi, and Mel Racenstein to the analysis of the structural MRI data and to the experimental volunteers who contributed their time to these studies.

