## Supplementary Materials for "Effects of cortical thickness, volume, and memory performance on age differences in neural reinstatement of scene information"

### Do regional structural metrics predict scene reinstatement?

Mirroring our approach in the methods section, we used a multiple regression approach to perform separate analyses with either PPA, MPA, or OPA reinstatement indices as the dependent variable. Cortical ROIs were defined using the Destrieux atlas (Destrieux et al. 2010) and extracted using Freesurfer. Regions that best matched with ROIs used to estimate the reinstatement indices were used: a combination of the parahippocampal and fusiform gyri (PHG/FG; Destrieux parcels 21 and 23) was used for the PPA, the ventral posterior cingulate cortex region (vPCC; Destrieux parcel 10) was used for the MPA, and a combination of the middle and superior occipital gyri (MOG/SOG; Destrieux parcels 19 and 20) was used for the OPA. The relevant structural metric (volume or thickness) of the remaining cortex (adjusted for total intracranial volume in the case of volumetric data) was added to any model whenever a regional metric was reliably associated with reinstatement. The rationale for this procedure was to ensure that any regional associations were indeed regionally specific.

**Regional cortical thickness as a predictor of scene reinstatement.** Analogously to the whole-brain volume measures used in our principal analyses, the initial regression models employed age group, study, regional thickness, and the interactions between age group and regional thickness and study and regional thickness as predictors of PPA, MPA, or OPA reinstatement. The interaction terms were non-significant in all cases for PPA and OPA reinstatement (PPA model: age × PHG/FG thickness: $p = .606$, study × PHG/FG thickness: $p = .109$; OPA model: age × MOG/SOG thickness: $p = .725$, study × MOG/SOG thickness: $p = .804$), and, therefore, these models were re-run after their exclusion. In the MPA model, there was a significant interaction between age and regional thickness (age × vPCC thickness: $p = .050$), while the study by regional thickness interaction was nonsignificant (study × vPCC thickness: $p = .166$). Because the significant interaction between age and vPCC thickness suggests that the relationship between neural reinstatement in the MPA and vPCC thickness may be age-dependent, we split the regression model between young and older adults. This split model revealed that vPCC thickness was not a significant predictor of reinstatement in this region in either age group (young adults: p = .070; older adults: p = .348). The outcomes from the reduced models are presented in Supplementary Table 1.

**Supplementary Table 1**. Outcome of linear regression analyses testing whether regional cortical thickness, age group, and study significantly predicted reinstatement in the PPA, MPA, and OPA.

| | | ***b*** | **SE *b*** | **β** | **t** | **Partial corr.** | **p-value** |
|---|---|---|---|---|---|---|---|
| **PPA** | PHG/FG thickness | .068 | .217 | .036 | .314 | .035 | .755 |
| **reinstatement** | Age group | -.197 | .072 | -.314 | -2.752 | -.298 | **.007*** |
| | Study | .016 | .068 | .026 | .241 | .027 | .810 |
| **MPA** | vPCC thickness | .080 | .127 | .070 | .631 | .071 | .530 |
| **reinstatement** | Age group | -.171 | .056 | -.335 | -3.040 | -.325 | **.003*** |
| | Study | -.037 | .054 | -.073 | -.689 | -.078 | .493 |
| **OPA** | MOG/SOG thickness | .166 | .194 | .110 | .855 | .096 | .395 |
| **reinstatement** | Age group | -.083 | .056 | -.185 | -1.482 | -.166 | .142 |
| | Study | -.087 | .051 | -.192 | -1.699 | -.189 | .093 |

Note: *b* = unstandardized coefficient; SE *b* = standard error of the unstandardized coefficient; β = standardized coefficient. Significant effects (p < .05) are in bold.

**Regional cortical volume as a predictor of scene reinstatement.** Again, mirroring our principal analyses, we set up initial regression models employing age group, study, regional volume, and the interactions between age group and regional volume and study and regional volume as predictors of reinstatement in the PPA, MPA, or OPA. Regional volume measures were adjusted for total intracranial volume (using the same process described in the methods section) to account for individual differences in brain size. The interaction terms for the models including regional volume were nonsignificant for the PPA (age × PHG/FG volume: *p* = .879; study × PHG/FG volume: *p* = .326), MPA (age × vPCC volume: *p* = .302; study × vPCC volume: *p* = .435), and OPA (age × MOG/SOG volume: *p* = .292; study × MOG/SOG volume: *p* = .692). Thus, all models were re-run excluding the interaction terms. The outcomes of these reduced models are presented in Supplementary Table 2.

**Supplementary Table 2**. Outcome of linear regression analyses testing whether regional cortical volume, age group, and study significantly predicted reinstatement in the PPA, MPA, and OPA.

| | | ***b*** | **SE *b*** | **β** | **t** | **Partial corr.** | **p-value** |
|---|---|---|---|---|---|---|---|
| **PPA** | PHG/FG volume | .055 | .036 | .173 | 1.521 | .170 | .132 |
| **reinstatement** | Age group | -.174 | .069 | -.278 | -2.524 | -.275 | **.014*** |
| | Study | -.011 | .069 | -.017 | -.159 | -.018 | .874 |
| **MPA** | vPCC volume | .020 | .028 | .076 | .698 | .079 | .487 |
| **reinstatement** | Age group | -.175 | .055 | -.343 | -3.206 | -.341 | **.002*** |
| | Study | -.043 | .055 | -.083 | -.777 | -.088 | .439 |
| **OPA** | MOG/SOG volume | .007 | .033 | .033 | .209 | .024 | .835 |
| **reinstatement** | Age group | -.099 | .061 | -.220 | -1.618 | -.180 | .110 |
| | Study | -.078 | .054 | -.173 | -1.458 | -.163 | .149 |

Note: *b* = unstandardized coefficient; SE *b* = standard error of the unstandardized coefficient; β = standardized coefficient. Significant effects ($p < .05$) are in bold.

**Do prefrontal structural metrics predict scene reinstatement?**

To investigate whether structural metrics derived from one or more regions of the prefrontal cortex might be the drivers of the whole-brain results, we used a similar multiple regression approach to that described above to determine whether thickness of volume of each of three prefrontal brain regions predicted reinstatement indices in the PPA, MPA, and OPA. Cortical ROIs for the superior frontal gyrus (SFG; Destrieux parcel 16), middle frontal gyrus (MFG; Destrieux parcel 15), and inferior frontal gyrus (IFG; Destrieux parcels 12, 13, and 14) were defined using the Destrieux atlas and extracted using Freesurfer.

**Prefrontal cortical thickness as a predictor of scene reinstatement.** The initial models employed the variables of age group, study, thickness of the given prefrontal region, and the interactions between age group and study with prefrontal regional thickness as predictors of PPA, MPA, or OPA reinstatement. The interaction terms were nonsignificant for all combinations of age and study with SFG, MFG, and IFG thickness in all models (Supplementary Table 3). Each model was therefore re-run excluding the interaction terms, and the outcomes of these reduced models are presented in Supplementary Tables 4–6.

**Supplementary Table 3.** Significance levels of interactions between age group and study with SFG, MFG, and IFG regional thickness in models predicting PPA, MPA, and OPA reinstatement.

| | **Interaction term** | **SFG** | **MFG** | **IFG** |
|---|---|---|---|---|
| **PPA reinstatement** | Age × thickness | .610 | .479 | .668 |
| | Study × thickness | .778 | .823 | .384 |
| **MPA reinstatement** | Age × thickness | .471 | .961 | .819 |
| | Study × thickness | .122 | .321 | .380 |
| **OPA reinstatement** | Age × thickness | .277 | .409 | .361 |
| | Study × thickness | .147 | .383 | .069 |

Note: Significant effects ($p < .05$) are in bold.

**Supplementary Table 4.** Outcome of linear regression analyses testing whether SFG thickness, age group, and study significantly predicted reinstatement in the PPA, MPA, and OPA.

| | | *b* | SE *b* | β | t | Partial corr. | p-value |
|---|---|---|---|---|---|---|---|
| **PPA** | SFG thickness | .044 | .202 | .033 | .218 | .025 | .828 |
| **reinstatement** | Age group | -.190 | .096 | -.303 | -1.987 | -.220 | **.050*** |
| | Study | .017 | .068 | .026 | .245 | .018 | .807 |
| **MPA** | SFG thickness | -.058 | .162 | -.054 | -.360 | -.041 | .720 |
| **reinstatement** | Age group | -.202 | .077 | -.395 | -2.628 | -.285 | **.010*** |
| | Study | -.036 | .054 | -.070 | -.667 | -.075 | .507 |
| **OPA** | SFG thickness | -.156 | .146 | -.164 | -1.069 | -.120 | .288 |
| **reinstatement** | Age group | -.159 | .069 | -.355 | -2.313 | -.253 | **.023*** |
| | Study | -.076 | .049 | -.167 | -1.552 | -.173 | .125 |

Note: *b* = unstandardized coefficient; SE *b* = standard error of the unstandardized coefficient; β = standardized coefficient. Significant effects ($p < .05$) are in bold.

**Supplementary Table 5.** Outcome of linear regression analyses testing whether MFG thickness, age group, and study significantly predicted reinstatement in the PPA, MPA, and OPA.

| | | ***b*** | **SE *b*** | **β** | **t** | **Partial corr.** | **p-value** |
|---|---|---|---|---|---|---|---|
| **PPA reinstatement** | MFG thickness | .096 | .196 | .071 | .491 | .055 | .625 |
| | Age group | -.175 | .091 | -.279 | -1.916 | -.212 | .059 |
| | Study | .015 | .068 | .024 | .226 | .026 | .822 |
| **MPA reinstatement** | MFG thickness | .056 | .157 | .051 | .359 | .041 | .721 |
| | Age group | -.164 | .073 | -.322 | -2.243 | -.246 | **.028*** |
| | Study | -.036 | .054 | -.070 | -.663 | -.075 | .509 |
| **OPA reinstatement** | MFG thickness | -.093 | .142 | -.097 | -.657 | -.074 | .513 |
| | Age group | -.136 | .066 | -.303 | -2.063 | -.227 | **.042*** |
| | Study | -.073 | .049 | -.161 | -1.491 | -.166 | .140 |

Note: *b* = unstandardized coefficient; SE *b* = standard error of the unstandardized coefficient; β = standardized coefficient. Significant effects ($p < .05$) are in bold.

**Supplementary Table 6.** Outcome of linear regression analyses testing whether IFG thickness, age group, and study significantly predicted reinstatement in the PPA, MPA, and OPA.

| | | ***b*** | **SE *b*** | **β** | **t** | **Partial corr.** | **p-value** |
|---|---|---|---|---|---|---|---|
| **PPA reinstatement** | IFG thickness | .216 | .231 | .141 | .933 | .105 | .354 |
| | Age group | -.142 | .095 | -.227 | -1.504 | -.168 | .137 |
| | Study | .020 | .067 | .032 | .300 | .034 | .765 |
| **MPA reinstatement** | IFG thickness | .212 | .185 | .171 | 1.147 | .129 | .255 |
| | Age group | -.120 | .076 | -.236 | -1.588 | -.177 | .116 |
| | Study | -.031 | .054 | -.061 | -.582 | -.066 | .562 |
| **OPA reinstatement** | IFG thickness | -.114 | .168 | -.104 | -.679 | -.077 | .499 |
| | Age group | -.140 | .069 | -.312 | -2.032 | -.224 | **.046*** |
| | Study | -.076 | .049 | -.168 | -1.549 | -.173 | .125 |

Note: *b* = unstandardized coefficient; SE *b* = standard error of the unstandardized coefficient; β = standardized coefficient. Significant effects ($p < .05$) are in bold.

**Prefrontal cortical volume as a predictor of scene reinstatement.** The initial models employed age group, study, volume of the given prefrontal region, and the interactions between age group and study with prefrontal regional thickness as predictors of PPA, MPA, or OPA reinstatement. The volume metrics were adjusted for total intracranial volume. The interaction terms were nonsignificant for all combinations of age and study with SFG, MFG, and IFG volume in all models (for a summary of each case, see Supplementary Table 7). Each model was therefore re-run excluding the interaction terms, and the outcomes of these reduced models are presented in Supplementary Tables 8–10.

**Supplementary Table 7.** Significance levels of interactions between age group and study with SFG, MFG, and IFG regional volume in models predicting PPA, MPA, and OPA reinstatement.

| | **Interaction term** | **SFG** | **MFG** | **IFG** |
|---|---|---|---|---|
| **PPA reinstatement** | Age × volume | .542 | .800 | .814 |
| | Study × volume | .203 | .425 | .177 |
| **MPA reinstatement** | Age × volume | .973 | .307 | .414 |
| | Study × volume | .821 | .683 | .812 |
| **OPA reinstatement** | Age × volume | .548 | .075 | .987 |
| | Study × volume | .237 | .154 | .111 |

Note: Significant effects (p < .05) are in bold.

**Supplementary Table 8.** Outcome of linear regression analyses testing whether SFG volume, age group, and study significantly predicted reinstatement in the PPA, MPA, and OPA.

| | | ***b*** | **SE *b*** | **β** | **t** | **Partial corr.** | **p-value** |
|---|---|---|---|---|---|---|---|
| **PPA reinstatement** | SFG volume | .110 | .047 | .346 | 2.356 | .258 | **.021*** |
| | Age group | -.052 | .092 | -.082 | -.563 | -.064 | .575 |
| | Study | -.006 | .066 | -.010 | -.095 | -.011 | .925 |
| **MPA reinstatement** | SFG volume | .101 | .037 | .390 | 2.718 | .294 | **.008*** |
| | Age group | -.042 | .073 | -.081 | -.571 | -.064 | .570 |
| | Study | -.056 | .052 | -.109 | -1.067 | -.120 | .289 |
| **OPA reinstatement** | SFG volume | .040 | .035 | .178 | 1.163 | .131 | .248 |
| | Age group | -.050 | .068 | -.112 | -.738 | -.083 | .463 |
| | Study | -.082 | .049 | -.181 | -1.667 | -.186 | .099 |

Note: *b* = unstandardized coefficient; SE *b* = standard error of the unstandardized coefficient; β = standardized coefficient. Significant effects (p < .05) are in bold.

**Supplementary Table 9.** Outcome of linear regression analyses testing whether MFG volume, age group, and study significantly predicted reinstatement in the PPA, MPA, and OPA.

| | | *b* | SE *b* | β | t | Partial corr. | p-value |
|---|---|---|---|---|---|---|---|
| **PPA reinstatement** | MFG volume | .094 | .052 | .296 | 1.823 | .202 | .072 |
| | Age group | -.065 | .101 | -.103 | -.639 | -.072 | .525 |
| | Study | -.008 | .068 | -.013 | -.125 | -.014 | .901 |
| **MPA reinstatement** | MFG volume | .067 | .042 | .260 | 1.620 | .180 | .109 |
| | Age group | -.082 | .082 | -.160 | -.999 | -.112 | .321 |
| | Study | -.053 | .054 | -.103 | -.973 | -.110 | .333 |
| **OPA reinstatement** | MFG volume | -.001 | .038 | -.006 | -.036 | -.004 | .972 |
| | Age group | -.109 | .075 | -.242 | -1.451 | -.162 | .151 |
| | Study | -.073 | .050 | -.162 | -1.466 | -.164 | .147 |

Note: *b* = unstandardized coefficient; SE *b* = standard error of the unstandardized coefficient; β = standardized coefficient. Significant effects ($p < .05$) are in bold.

**Supplementary Table 10.** Outcome of linear regression analyses testing whether IFG volume, age group, and study significantly predicted reinstatement in the PPA, MPA, and OPA.

| | | *b* | SE *b* | β | t | Partial corr. | p-value |
|---|---|---|---|---|---|---|---|
| **PPA reinstatement** | IFG volume | .098 | .047 | .310 | 2.082 | .229 | **.041*** |
| | Age group | -.068 | .093 | -.108 | -.727 | -.082 | .469 |
| | Study | -.006 | .067 | -.010 | -.097 | -.011 | .923 |
| **MPA reinstatement** | IFG volume | .046 | .039 | .178 | 1.191 | .134 | .237 |
| | Age group | -.118 | .076 | -.231 | -1.551 | -.173 | .125 |
| | Study | -.046 | .055 | -.089 | -.844 | -.095 | .401 |
| **OPA reinstatement** | IFG volume | .021 | .035 | .094 | .608 | .069 | .545 |
| | Age group | -.077 | .069 | -.171 | -1.114 | -.125 | .269 |
| | Study | -.079 | .050 | -.174 | -1.585 | -.177 | .117 |

Note: *b* = unstandardized coefficient; SE *b* = standard error of the unstandardized coefficient; β = standardized coefficient. Significant effects ($p < .05$) are in bold.

Because SVG volume was a significant predictor of PPA and MPA reinstatement, and IFG volume significantly predicted PPA reinstatement, the volume of the remaining cortex (adjusted for total intracranial volume) was added to each of the respective models to assess whether the associations were regionally specific. As is apparent from Supplementary Table 11, neither prefrontal regional metric accounted for a significant amount of the variance in reinstatement after controlling for the volume of the remaining cortex.

**Supplementary Table 11.** Outcome of linear regression analyses testing whether SFG or IFG volume, the volume of the remaining cortex, age group, and study significantly predicted reinstatement in cases where SFG or IFG volume significantly predicted reinstatement before accounting for the volume of the remaining cortex.

| | | ***b*** | **SE *b*** | **β** | **t** | **Partial corr.** | **p-value** |
|---|---|---|---|---|---|---|---|
| **PPA reinstatement** | SFG volume | .010 | .065 | .032 | .154 | .018 | .878 |
| | Volume of remaining cortex | .149 | .070 | .468 | 2.134 | .236 | **.036*** |
| | Age group | .021 | .096 | .033 | .217 | .025 | .829 |
| | Study | -.053 | .068 | -.084 | -.776 | -.088 | .440 |
| **MPA reinstatement** | SFG volume | .055 | .053 | .213 | 1.042 | .118 | .301 |
| | Volume of remaining cortex | .068 | .056 | .262 | 1.203 | .136 | .233 |
| | Age group | -.008 | .078 | -.017 | -.109 | -.012 | .913 |
| | Study | -.077 | .055 | -.150 | -1.399 | -.157 | .166 |
| **PPA reinstatement** | IFG volume | -.015 | .065 | -.046 | -.224 | -.026 | .823 |
| | Volume of remaining cortex | .169 | .070 | .531 | 2.424 | .266 | **.018*** |
| | Age group | .016 | .096 | .026 | .166 | .019 | .869 |
| | Study | -.055 | .068 | -.086 | -.807 | -.092 | .422 |

Note: *b* = unstandardized coefficient; SE *b* = standard error of the unstandardized coefficient; β = standardized coefficient. Significant effects ($p < .05$) are in bold.

**ROI intercorrelations and associations with whole-brain cortical thickness and volume**

We employed partial correlations, controlling for age group and study, to estimate the correlations between the mean thickness of each ROI, and of the mean thickness of each ROI with mean whole-brain cortical thickness. The outcomes of these correlations are presented in Supplementary Table 12.

**Supplementary Table 12.** Correlation matrix showing the partial correlations between the mean thickness of each ROI with each other ROI and whole-brain cortical thickness, controlling for age group and study.

| | **PHG/FG** | **vPCC** | **MOG/SOG** | **SFG** | **MFG** | **IFG** | **Whole-brain cortical thickness** |
|---|---|---|---|---|---|---|---|
| **PHG/FG** | - | **.276 (.013)*** | **.412 (<.001)*** | **.464 (<.001)*** | **.462 (<.001)*** | **.422 (<.001)*** | **.592 (<.001)*** |
| **vPCC** | | - | **.274 (.014)*** | .180 (.109) | .102 (.368) | .204 (.069) | **.304 (.006)*** |
| **MOG/SOG** | | | - | **.544 (<.001)*** | **.595 (<.001)*** | **.518 (<.001)*** | **.713 (<.001)*** |
| **SFG** | | | | - | **.874 (<.001)*** | **.594 (<.001)*** | **.772 (<.001)*** |
| **MFG** | | | | | - | **.707 (<.001)*** | **.808 (<.001)*** |
| **IFG** | | | | | | - | **.792 (<.001)*** |
| **Whole-brain cortical thickness** | | | | | | | - |

Note: Strength and direction of correlations are shown with significance levels in parentheses. Significant correlations ($p < .05$) are shown in bold.

Next, we ran another series of partial correlations controlling for age group and study to identify the correlations between the volumes of each ROI, and of the volumes of each ROI with total cortical volume. The outcomes of these correlations are presented in Supplementary Table 13.

**Supplementary Table 13.** Correlation matrix showing the correlations between the volumes of each ROI with each other ROI and whole-brain cortical volume, controlling for age group and study.

| | PHG/FG | vPCC | MOG/SOG | SFG | MFG | IFG | Whole-brain cortical volume |
|---|---|---|---|---|---|---|---|
| **PHG/FG** | - | .156 (.166) | **.327 (.003)*** | **.420 (<.001)*** | **.272 (.015)*** | **.385 (<.001)*** | **.531 (<.001)** |
| **vPCC** | | - | **.384 (<.001)*** | **.221 (.049)*** | **.257 (.022)*** | .143 (.205) | **.304 (.006)*** |
| **MOG/SOG** | | | - | **.233 (.038)*** | **.400 (<.001)*** | **.344 (.002)*** | **.492 (<.001)*** |
| **SFG** | | | | - | **.613 (<.001)*** | **.627 (<.001)*** | **.764 (<.001)*** |
| **MFG** | | | | | - | **.541 (<.001)*** | **.684 (<.001)*** |
| **IFG** | | | | | | - | **.737 (<.001)*** |
| **Whole-brain cortical volume** | | | | | | | - |

Note: Strength and direction of correlations are shown with significance levels in parentheses. Significant correlations ($p < .05$) are shown in bold.
